# Tracking evolutionary trends towards increasing complexity: a case study in Cyanobacteria

**DOI:** 10.1101/2020.01.29.924464

**Authors:** Andrés Moya, José L. Oliver, Miguel Verdú, Luis Delaye, Vicente Arnau, Pedro Bernaola-Galván, Rebeca de la Fuente, Wladimiro Díaz, Cristina Gómez-Martín, Francisco M. González, Amparo Latorre, Ricardo Lebrón, Ramón Román-Roldán

**Author notes:** These authors contributed equally (José L. Oliver, Miguel Verdú, Luis Delaye). Corresponding author: Andrés Moya.

## Abstract

Progressive evolution, the tendency towards increasing complexity, is a controversial issue in Biology, whose resolution requires a proper measurement of complexity. Genomes are the best entities to address this challenge, as they record the history and information gaining of organisms in their ongoing biotic and environmental interactions. Using six metrics of genome complexity, none of which is primarily associated to biological function, we measure genome complexity in 91 genomes from the phylum Cyanobacteria. Several phylogenetic analyses reveal the existence of progressive evolution towards higher genome complexity: 1) all the metrics detect strong phylogenetic signals; 2) ridge regressions detect positive trends towards higher complexity; and 3) classical proofs for progressive evolution (the minimum, the ancestor-descendent and the sub-clade tests), show that some of these positive trends are driven, being mainly due to natural selection. These findings support the existence of progressive genome evolution in this ancient and diverse group of organisms.

## Introduction

Treatises on biological evolution reflect a conflict between the relative roles played by contingency and necessity (Moya, 2014). An important tradition in Evolutionary Biology, based on a large amount of empirical evidence, considers that contingency marks the dynamics of evolution in a way that makes it unpredictable. The trend towards the appearance of increasing complexity falls within the frame of contingent evolution insofar as it is inevitable given that, passively, we can expect that sooner or later more complex entities will evolve from the original, simpler entities. This is what Gould (1996) calls the passive tendency towards complexity marked by the minimum initial complexity wall.

Assuming there is an evolutionary trend towards greater complexity, a fundamental question is whether we can prove the existence of a metric accounting for it (McShea and Brandon, 2010; Day, 2012). We first conjecture that it is in the genomes where we can find evidence of such metrics, which may eventually increase over evolutionary time. Genomes record the history and information gained by organisms during their interaction with environmental and biotic factors over time (Adami, 2002, 2016; Krakauer, 2011). However, genome parameters such as genome size, number of genes, gene components (i.e., introns, exons), etc., are insufficient to show any evolutionary trend. We speculate that this is probably because they only partially capture the abovementioned history and information gain (Lieberman-Aiden *et al*., 2009; Dekker *et al*., 2017). Our second conjecture is that the best way to measure genome complexity is by using metrics that are not primarily associated to biological function.

The currently available metrics applied to genomes are very broad and not all of them appropriately capture the information gained by the genome during its evolutionary history (Zurek, 1990; Adami, 2002). Algorithmic complexity (Chaitin, 1977; Li and Vitányi, 2008) is inconveniently maximum for randomness. The effective complexity of Gell-Mann and Lloyd (1996) is recommended for collections or ensembles of sequences, but in many cases, such as in genome sequences, it is not clear how to define an appropriate ensemble. Likewise, those based on mutual information, i.e. statistical dependence (Grassberger, 1986; Adami and Cerf, 2000) also quantify the complexity of sequence ensembles generated by a given source rather than the complexity of a single sequence. Here we consider metrics based on an individual entity (genome). Interestingly, four of them consider the number of different parts composing the genome and the irregularity of their distribution, thus extrapolating to DNA sequences the operational definition of McShea (1993), which measures biological complexity as the degree to which the parts of a morphological structure differ from each other. Two additional metrics are based on *k-mer* statistics. A key point is that none of the six metrics we used to measure genome complexity considers biological function.

A first group of metrics are based on the Sequence Compositional Complexity derived from a four-symbol DNA sequence (*SCC*) or from the binary sequences resulting from grouping the four nucleotides into S(C,G) vs. W(A,T) or R(A,G) vs. Y (T,C), or K(A,C) vs. M(T,G), thus obtaining *SCC_SW_*, *SCC_RY_* and *SCC_KM_* metrics, respectively (Román-Roldán *et al*., 1998). These four metrics increase with the number of parts (i.e. compositional domains) found in a genome sequence by a segmentation algorithm, and both the length and compositional differences among them.

The fifth metric we used is the Biobit (*BB*), a metric based on the difference between the maximum entropy for a *k*-*mer* of a random genome of same length as the genome under consideration and the entropy of that genome for such a *k*-mer (Bonnici and Manca, 2016). Lastly, we used the Genomic Signature (*GS*), also a *k*-*mer*-based metric, which is the value corresponding to the *k*-*mer* that maximizes the difference between observed and expected equi-frequent classes of *mers*. *GS* is based on the relative abundances of short oligonucleotides (Karlin and Ladunga, 1994) and chaos game representation applied to genomes (Jeffrey, 1990; Almeida *et al*., 2001).

As already stated, the four *SCCs* are the only metrics using genome segmentation (i.e., the partition of the genome into non-overlapping fragments of varying lengths and with homogeneous composition). These metrics parallel the concept of ‘pure complexity’ of McShea and Brandon (2010), where complexity is more closely related with the number of part types of a living being than with the number of functions. The other two metrics, based on distribution of *k*-*mers*, do not use genome partitioning.

We test the above-mentioned metrics by analyzing the genome evolution of an ancient and diverse group of organisms: the Cyanobacteria phylum. These microorganisms played a fundamental role in the evolution of life on Earth. The fossil record shows that they were here 2.0 billion years ago (Bya) and molecular clock analyses indicate that the phylum originated over 2.5 Bya (Sergeev *et al*., 2002; Schirrmeister *et al.,* 2013). By releasing oxygen through photosynthesis, Cyanobacteria caused the Great Oxidation Event about 2.3 Bya and changed the history of life on Earth (Bekker *et al.,* 2004). The oxidation of the environment allowed the evolution of complex multicellular life forms (Hedges *et al*., 2004), leading to the origin of eukaryotic crown groups including plants and animals (Knoll, 2014). As it is well known, Cyanobacteria originated plastids through symbiosis with ancient eukaryotes (Sagan, 1967).

Cyanobacteria are morphologically diverse. The group was traditionally classified into five subsections according to several biological criteria (Rippka *et al*., 1979; Rippka, 1988). These subsections of Cyanobacteria are not monophyletic (Schirrmeister *et al.,* 2013; Dagan *et al.,* 2013). More recent classification schemes using phylogenetic analysis from 31 conserved protein sequences divide Cyanobacteria into nine different groups (Komárek *et al*., 2014). These are: Gleobacterales, Synechococcales, Oscillatoriales, Chroococcales, Pleurocapsales, Spirulinales, Rubidibacter/Halothece, Chroococcidiopsidales and Nostocales. Not all these groups are monophyletic. Clearly, taxonomy and evolution of Cyanobacteria is an active area of research and this classification is likely to change in the near future.

As a proof of concept, in the present work we test whether there is a statistically and phylogenetically supported driven tendency towards increasing genome complexity as reflected by the metrics of genome complexity and/or by genome standard parameters (genome size, %GC and number of genes) in the evolution of Cyanobacteria.

## Results

### Complexity measures throughout Cyanobacteria phylogeny

The values of the four *SCCs*, *BB* and *GS* metrics as well as three standard genome parameters (Genome size, %GC and No. of genes) (see material and methods) for 91 Cyanobacterial genomes are reported in Supplementary file 1. Figure 1 shows a maximum likelihood phylogeny of Cyanobacteria whose branch lengths are proportional to the number of amino acid substitutions (see material and methods). The phylogeny is in general agreement with previous analysis (Dagan *et al*., 2013; Schirrmeister *et al*., 2013; Komárek *et al*., 2014).

**Figure 1.**
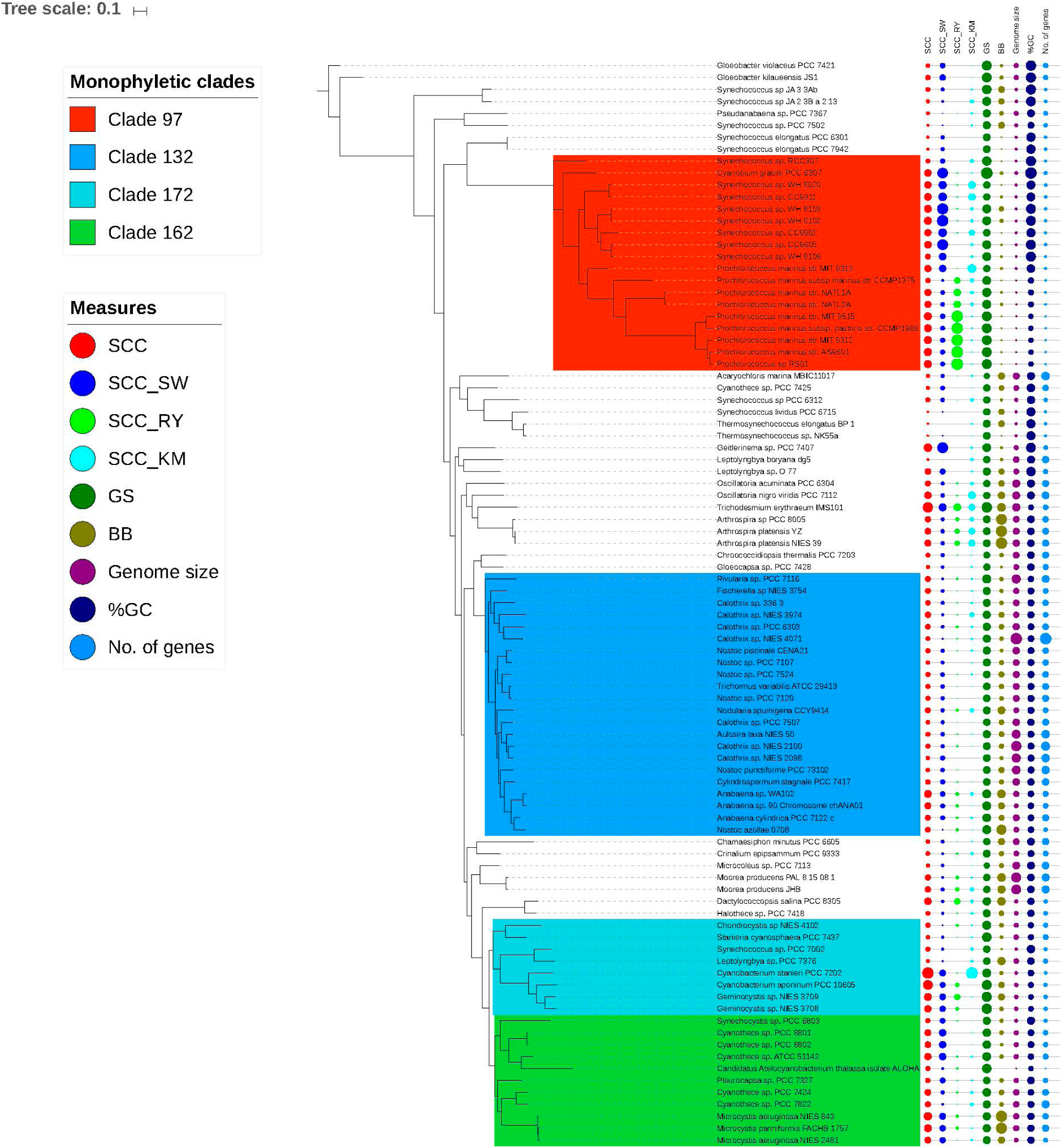
Phylogeny of Cyanobacteria with the metrics of complexity and genome parameters for each chosen genome. Values of metrics and parameters are proportional to circle size and were standardized to have a mean of zero and variance of one. The four colored boxes represent four monophyletic sub-clades that were used to test evidence of a driven trend for each sub-clade.

### Phylogenetic signal

All metrics of complexity and genome parameters showed a significant phylogenetic signal (Table 1), indicating that for all cases genomes of closely related cyanobacterial species tend to resemble more than two randomly selected genomes (Figure 1). However, the magnitude of the phylogenetic signal differs across metrics and parameters, with %GC and *GS* showing the highest values, which probably reflects different forces shaping the evolution of all these metrics and parameters (see Discussion).

**Table 1.** Phylogenetic signals (*K*) of the six metrics of genome complexity and the three genome parameters.

| Genome parameters and metrics of genome complexity | $K$ | Probability, $p$ |
| --- | --- | --- |
| $SSC$ | 0.34 | 0.001 |
| $SCC_{RY}$ | 0.66 | 0.001 |
| $SCC_{SW}$ | 0.32 | 0.001 |
| $SCC_{KM}$ | 0.26 | 0.001 |
| $BB$ | 0.15 | 0.001 |
| $GS$ | 1.00 | 0.001 |
| Genome size | 0.46 | 0.001 |
| %GC | 3.96 | 0.001 |
| No. of genes | 0.31 | 0.001 |

### Phylogenetic correlations

To gain a better understanding of the metrics, after correction of the phylogenetic signals, we evaluated how they correlate with each other and, particularly, with the genomic parameters (see Table 2). It is worth noticing that two metrics in particular, *SCC* and *SCC_RY_* correlate with other metrics and parameters (six correlations each one), accounting for 43% of all significant correlations.

**Table 2.** Phylogenetic Pearson correlations (*r*) among metrics of genome complexity and genome parameters. Statistical significance was corrected by false discovery rates to control for multiple testing.

| | $SCC$ | $SCC_{SW}$ | $SCC_{RY}$ | $SCC_{KM}$ | $BB$ | $GS$ | Genome<br>Size | %GC |
| --- | --- | --- | --- | --- | --- | --- | --- | --- |
| $SCC_{SW}$ | 0.66<sup>***</sup> | | | | | | | |
| $SCC_{RY}$ | 0.52<sup>***</sup> | 0.09<sup>ns</sup> | | | | | | |
| $SCC_{KM}$ | 0.30<sup>**</sup> | 0.09<sup>ns</sup> | 0.03<sup>ns</sup> | | | | | |
| $BB$ | 0.38<sup>***</sup> | -0.02<sup>ns</sup> | 0.53<sup>***</sup> | -0.04<sup>ns</sup> | | | | |
| $GS$ | 0.34<sup>***</sup> | 0.20<sup>ns</sup> | 0.41<sup>***</sup> | -0.20<sup>ns</sup> | 0.19<sup>ns</sup> | | | |
| Genome Size | 0.22<sup>*</sup> | 0.10<sup>ns</sup> | 0.31<sup>**</sup> | -0.05<sup>ns</sup> | 0.32<sup>**</sup> | 0.001<sup>ns</sup> |  |  |
| %GC | -0.06<sup>ns</sup> | 0.26<sup>*</sup> | -0.38<sup>***</sup> | -0.11<sup>ns</sup> | -0.09<sup>ns</sup> | -0.1<sup>ns</sup> | -0.11<sup>ns</sup> |  |
| No. of genes | 0.12<sup>ns</sup> | 0.07<sup>ns</sup> | 0.26<sup>*</sup> | -0.09<sup>ns</sup> | 0.26<sup>*</sup> | -0.09<sup>ns</sup> | 0.86<sup>***</sup> | -0.09<sup>ns</sup> |

\*\*\* $p < 0.001$ ; \*\* $0.001 < p < 0.01$ ; \* $0.01 < p < 0.05$ ; <sup>ns</sup> $p > 0.05$

### Ridge regression of complexity metrics *versus* age

Using ridge regression of genome complexity metrics and genome parameters *versus* age (distance from the root), we have studied whether there is evidence of evolutionary trends, having detected interesting patterns (Figure 2, panels A and B). Of the complexity metrics, four out of six show a statistically significant positive trend (*SCC, SSC_SW_*, *SCC_RY_* and *GS*), indicating that complexity has increased with evolutionary time. In contrast, *SCC_KM_* shows no trend and *BB* reveals a significant negative trend. Interestingly genome parameters, on the other hand, show no evidence of any evolutionary trend.

**Figure 2.**
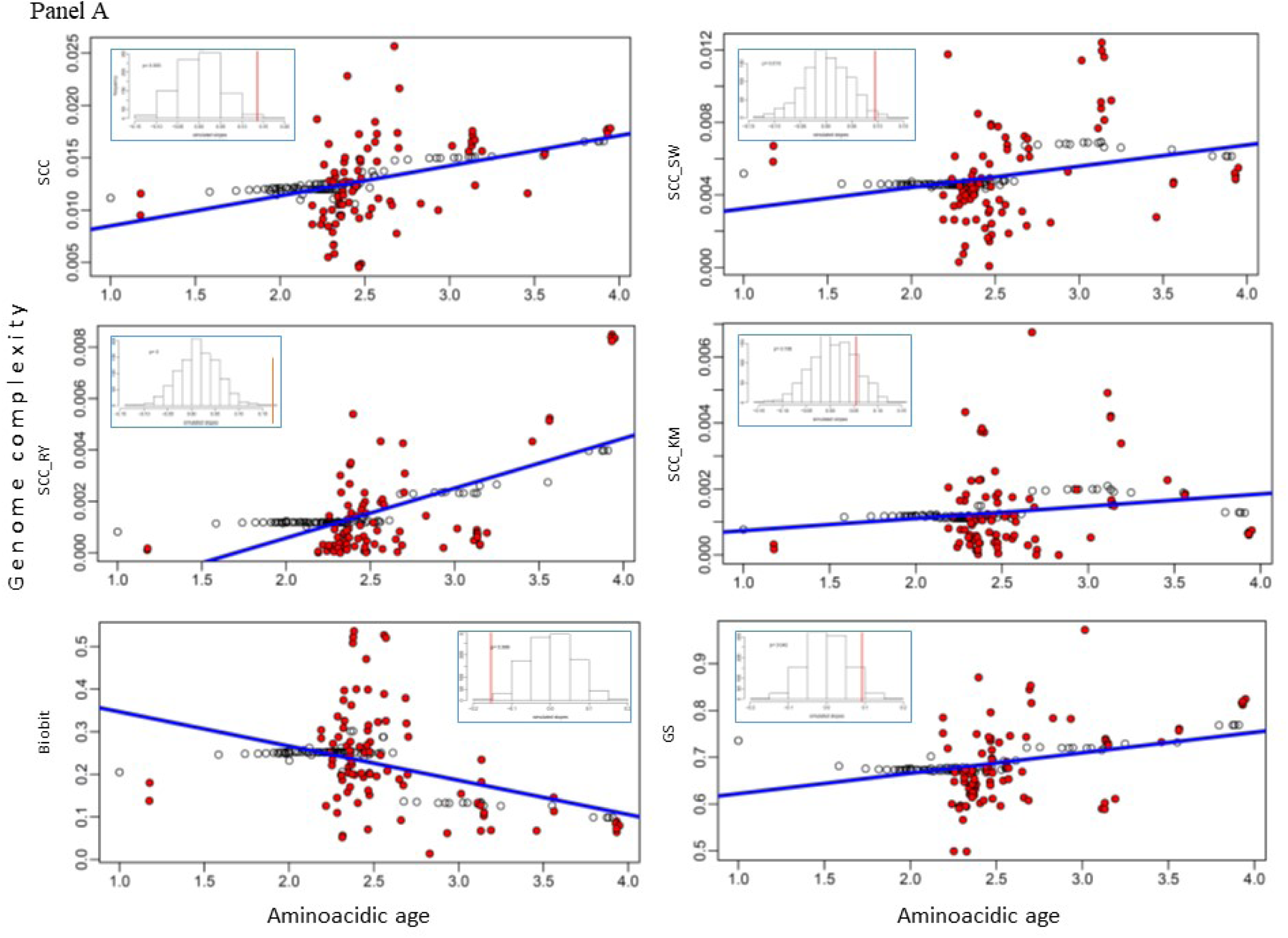
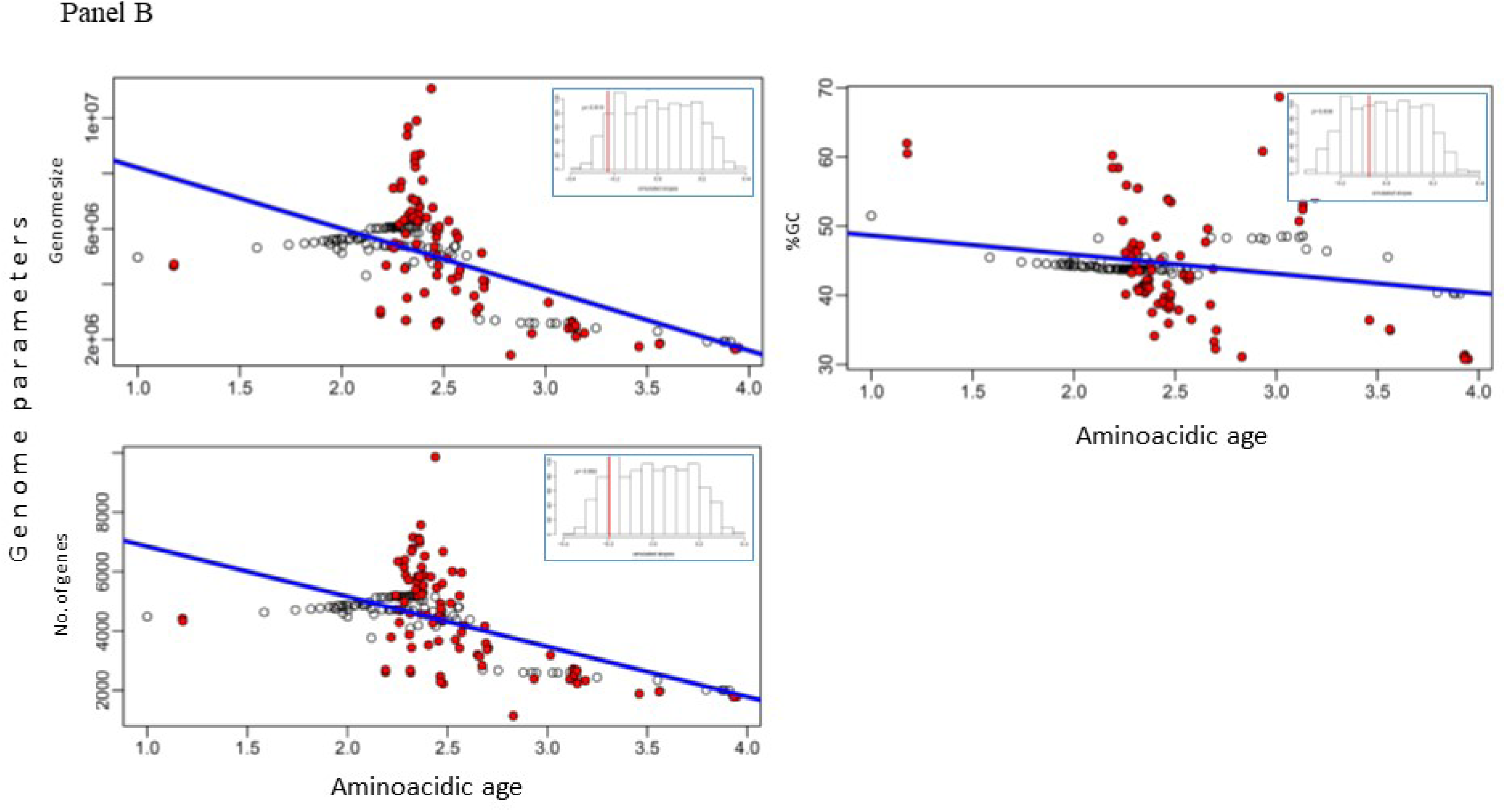
Phylogenetic trends of genomic complexity metrics (panel A) and standard genome parameters (panel B). The estimated genomic value for each tip (red circles) or node (white circles) in the phylogenetic tree is regressed against its evolutionary age (i.e., distance from the root). The statistical significance of the regression (blue lines) is tested against 10,000 slopes obtained under simulated Brownian evolution. The upper plot shows the frequency distribution of the 10,000 simulated slopes and the red vertical line shows the estimated slope. The slopes and their *p*-values are shown in Supplementary file 4.

### Driven trends in Cyanobacteria

To unravel whether the positive trends are passive or driven we have applied three types of proofs, called the minimum, the ancestor-descendent and the subclade proofs, respectively (McShea, 1994; see also McShea and Brandon, 2010). These proofs are well known in Paleontology and Evolutionary Biology and, to the best of our knowledge, this is the first time they have been applied to genome evolutionary analyses. To gain a better understanding of the positive trends we have also applied those proofs for comparative purposes to the metrics and genome parameters that do not show evidence of such a positive evolutionary trend.

#### Minimum proof

Regarding minimum proof, we have applied three types of tests. The first one is the skewness of the entire phylum. In this respect, we observed that the skewness of all metrics (except *SCC* and GC content) exhibit significant and positive skewness (D’Agostino-Pearson test, *n* = 91; see Table 3). This result supports a left wall for these metrics and parameters, which is compatible with either a passive or a driven trend. On the other hand, it is expected that if the minimum value of a given metric or parameter delimiting the left wall increases with evolutionary time, then the trend will probably be driven. To evaluate this, we considered as the minimum the estimated value of the most basal clade, *xb*, for each metric/parameter (see Figure 1). However, as it can be observed (Figure 3), there are lower or higher values than the one corresponding to the basal clade for any metric/parameter. A deeper study of the distribution of lower and higher values with respect to *xb* could give us evidence of the putative existence of a driven trend. To test this (second test of the minimum proof) we measure |*xd-xb*|, the absolute difference between descendants’ clades and the most basal clade. Table 3 shows whether there is a statistical difference (Chi-square test) between those clades (179 in total) that are higher or lower than the basal clade, *xb*. As it can be observed, all the tests are significant with four metrics (*SCC*, *SCC_RY_*, *SCC_KM_* and *BB*) and two parameters (Genome size and No. of genes) showing a significant increase in the metric/parameter with respect to the corresponding basal values. In contrast, two metrics (*SCC_SW_* and *GS*) and one parameter (%GC) present a significant decrease. We have also tested by means of a Student’s t-test (third test of the minimum proof) whether there is a statistical difference between the average value of the absolute difference (|*xd-xb*|) of a give metric or parameter higher or lower than *xb*. In this case, it can be observed (see Table 3) that three metrics (*SCC_SW_*, *SCC_RY_* and *SCC_KM_*) show a statistical difference in favor of a higher value than *xb* and one metric (*GS*) and the three parameters (Genome size, %GC content and No. of genes) present a statistical difference in favor of a lower value than *xb*.

**Figure 3.**
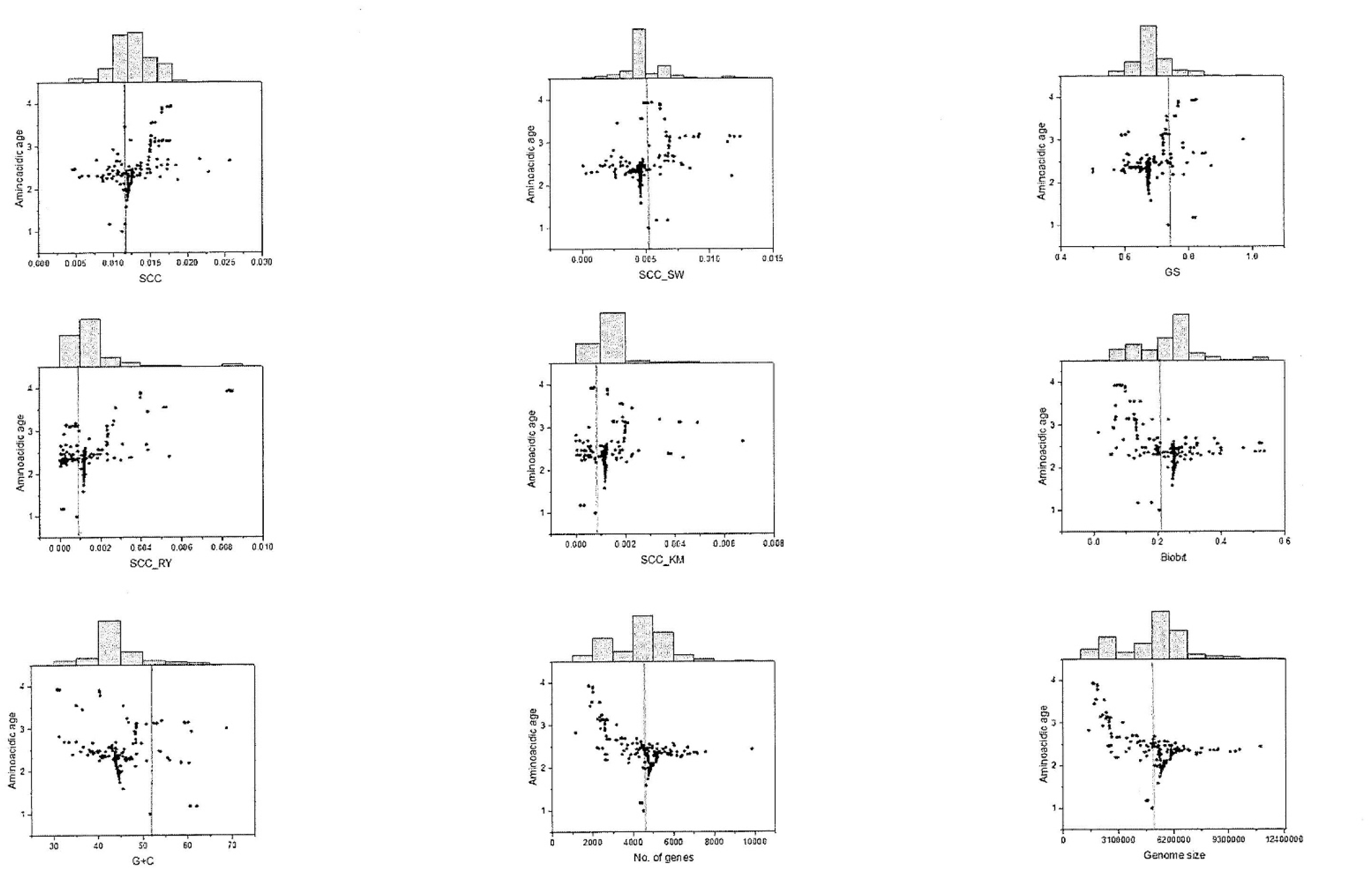
Distribution of metrics and parameters according to amino-acidic age. The interior dashed line corresponds to the value of the basal clade, *xb*. The histograms that appear above each figure correspond to the number of accumulated values of metrics and parameters (regardless of the age) ranging from lower (left) to higher (right) values than *xb*.

**Table 3.**
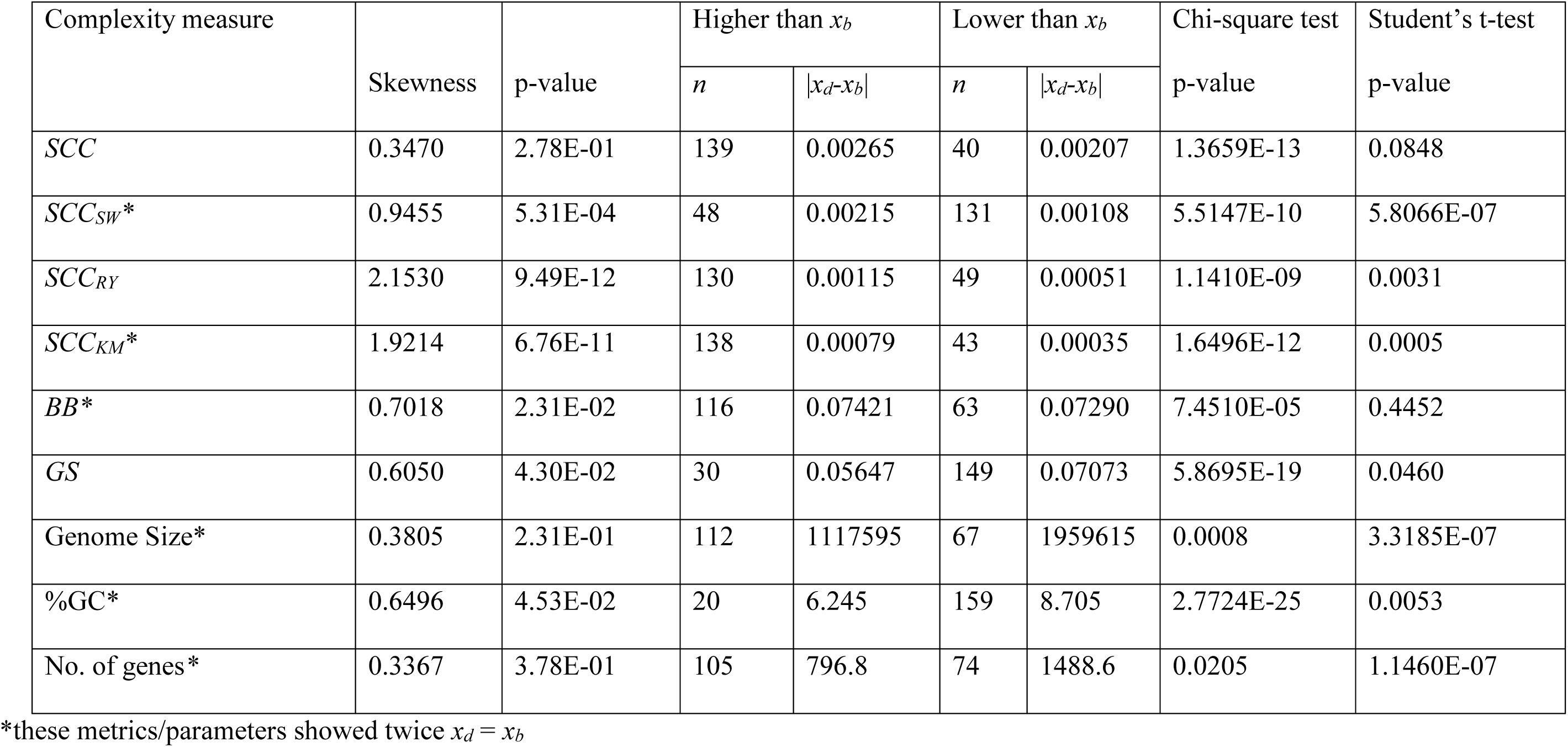
Proofs of the minimum. D’Agostino-Pearson test of skewness for the entire Phylum (n = 91) and number (*n*) of times that the metric/parameter of a given derived internal or terminal node (*xd*) is higher or lower than the basal node (*xb*) (Chi-square test), as well as the average absolute difference (|*xd-xb*|, Student’s t-test) between nodes that are higher or lower than *xb*.

#### The ancestor-descendent proof

According to Gould (1996) the ancestor-descendent proof is the most appropriate one to discover whether positive trends are passive or driven. McSchea (1994) indicates that in a passive system, increases and decreases should be the same, whereas in a driven trend the number of increases should occur more often. To test this, we tabulated the derived clades for all possible nodes and whether they present a higher, lower or equal value of the metric/parameter than the ancestral clade corresponding to each node. In order to avoid bias due to proximity to the putative left wall, McSchea (1994) recommends applying the test only to those clades where both ancestor and descendent are higher than the average value of the metric/parameter. As it can be observed (Table 4) this exigent test shows that metrics *SCC* and *GS* and the three genome parameters are in favor of a driven trend. A good visualization of the ancestor-descendant proof on the phylogeny of the Cyanobacteria for each metric/parameter has been obtained by estimating the values of internal nodes using a maximum likelihood function and interpolating the value along each edge. Figure 4 shows the mapping corresponding to the *SCC* metric where the driven positive trend of this metric can be clearly appreciated (see Supplementary file 2 for the mapping of the rest of metrics/parameters).

**Figure 4.**
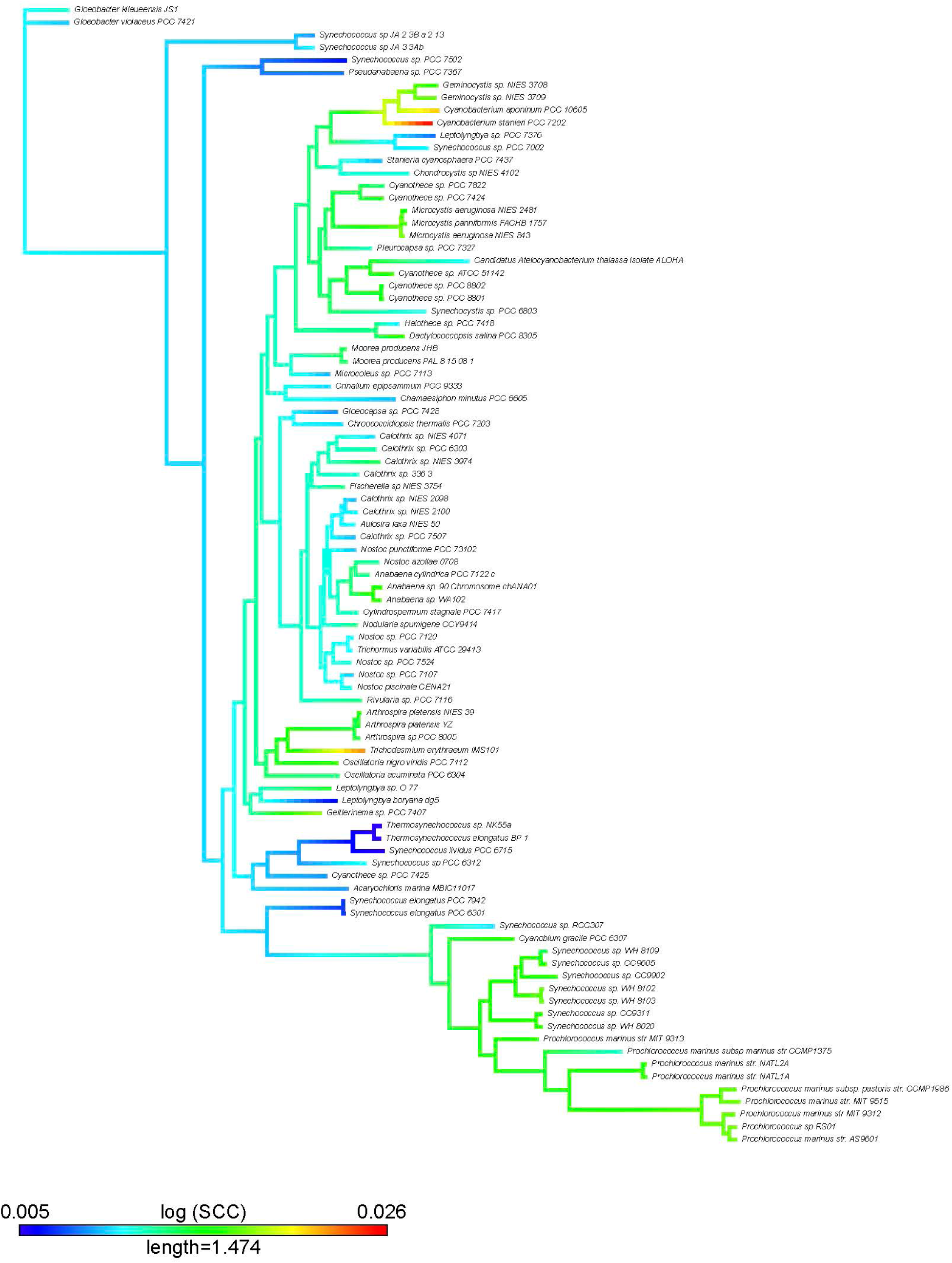
Mapping of the *SCC* complexity metric on the Cyanobacteria tree. We used two functions (*contMap* and *fastAnc*) from the *phytools* R package (Revell, 2012). The *contMap* R function allows plotting a tree with a mapped continuous character, such as any of our complexity measures. Mapping is accomplished by estimating states at internal nodes using maximum likelihood with the function *fastAnc* and interpolating the states along each edge using equation 2 of Felsenstein (1985).

**Table 4.** Ancestor-descendent proof. For any complexity measure/genome parameter we test whether the derived clades present higher or lower values than the corresponding ancestral clade for any node.

| Complexity measure | Derived nodes with a higher value than the ancestor of a given clade | Derived nodes with a lower value than the ancestor of a given clade | Fisher exact test $p$-value |
| --- | --- | --- | --- |
| *SCC* | 36 | 2 | 0.0001 |
| *SCC<sub>SW</sub>* | 19 | 9 | 0.2772 |
| *SCC<sub>RY</sub>* | 15 | 5 | 0.1908 |
| *SCC<sub>KM</sub>* | 15 | 15 | 1.0000 |
| *BB* | 58 | 32 | 0.0703 |
| *GS* | 33 | 5 | 0.0011 |
| Genome Size | 68 | 36 | 0.0018 |
| %GC | 68 | 8 | 0.0350 |
| No. of genes | 38 | 32 | 0.0143 |

#### The sub-clade proof

The final proof applied is the sub-clade proof. According to McSchea (1993) if the parent distribution is skewed (see histograms of Figure 3 and Table 3) and the mean skew of a sub-clade drawn from the right tail is also skewed, the system is probably driven. For this proof we have applied two types of test. First, we tested whether the trend observed at Phylum level is also observed in four selected monophyletic clades and second, we have also applied the skewness test proposed by McSchea (1993) properly to either the entire phylum (results given in Table 3) and to the chosen sub-clades. We have chosen four monophyletic clades formed by clusters 97, 132, 162 and 172 that harbor 18, 22, 11 and 8 species, respectively (four color boxes in Figure 1 and Supplementary file 3). Clade 97 is formed by Synechococcus, Prochlorococcus and Cyanobium; clade 132 corresponds to Nostocales (subsections IV and V, Komarek *et al*., 2012); clade 162 contains Cyanothece and Microcystis; and clade 172, among others, contains Geminocystis and Cyanobacterium. The most relevant result found was that some metrics of genome complexity show statistically significant positive trends (*SCC*, *SCC_RY_* and *GS*) and some others show negative trends (*SCC_SW_* and *SCC_KM_*), whereas the genomes parameters do not show any positive trends (see Supplementary files 4 and 5). Thus, we keep *SCC*, *SCC_RY_* and *GS* as the metrics showing positive trends at both levels of phylogenetic resolution.

Regarding the second test for sub-clade proof, we followed the criteria given by McShea, (1994) whereby the monophyletic sub-clade drawn from the right tail of the entire distribution should have a statistically significant average higher value than the one corresponding to the entire phylum. Regarding the skewness of the phylum (Table 3) we observe that all metrics (except *SCC* and %GC) exhibit significant and positive skewness. However, this test of skewness cannot be applied to the four chosen monophyletic sub-clades either because a) the average value (median) of a given metric/parameter for each sub-clade was lower than the median of the phylum (16 cases out of 36) or, b) there was no statistical evidence (the remaining 20 cases) of a higher median (Mood’s median test) of a given metric/parameter for each sub-clade than the median of the entire phylum (see Supplementary file 6).

In summary, the overall results obtained in relation to the evidence found for a trend in a given metric or parameter (i.e., the phylogenetic signal, the number of significant correlations against the rest of metrics/parameters, as well as whether the trend is driven or not (see Table 5)) show that *SCC*, *SCC_RY_* and to a lower extent *GS* present the highest scores, and can thus be considered metrics evidencing progressive evolution of Cyanobacteria.

**Table 5.** Summary of the results for each complexity metric and genome parameter. *K* is the phylogenetic signal. The signs “+”, “-“ or “0” indicate the existence of a positive, negative or no statistical evidence, respectively, for the corresponding test: the general trend, the driven trend, the three types of test of the minimum (i.e., skewness, Chi-square test and Student’s t-test), the ancestor-descendant test and the trend in the case of the four sub-clades.

| Complexity metric / parameter | $K$ | Number of significant correlations | General trend | Driven trend | | | | | | |
| --- | --- | --- | --- | --- | --- | --- | --- | --- | --- | --- |
|  |  |  |  | Minimum test |  |  | Ancestor-descendant test | Trend in the four sub-clades |  |  |
|  |  |  |  | Skewness | Chi-square test | Student’s t-test |  | + | - | 0 |
| $SCC$ | + | 6 | + | 0 | + | 0 | + | 2 | 0 | 2 |
| $SCC_{SW}$ | + | 2 | + | + | - | + | 0 | 1 | 2 | 1 |
| $SCC_{RY}$ | + | 6 | + | + | + | + | 0 | 2 | 0 | 2 |
| $SCC_{KM}$ | + | 1 | 0 | + | + | + | 0 | 0 | 2 | 2 |
| $BB$ | + | 4 | - | + | + | 0 | 0 | 0 | 1 | 3 |
| $GS$ | + | 2 | + | + | - | - | + | 3 | 0 | 1 |
| Genome Size | + | 4 | - | + | + | - | + | 0 | 2 | 2 |
| %GC | + | 2 | - | 0 | - | - | + | 0 | 2 | 2 |
| No. of genes | + | 3 | - | + | + | - | + | 0 | 2 | 2 |

## Discussion

Genomes probably provide the best record of the biological history of species, not only do they enable us to reconstruct their phylogenetic relationships but they also contain information gained from their continuous biotic and environmental interactions over time (Adami, 2002; Krakauer, 2011). This information is an elusive but crucial component of the genome, whose study as a whole deserves deeper attention because it holds clues to answer many biological questions, particularly those of an evolutionary nature. The genome has distinct layers of information encoded in DNA sequences (Dekker *et al*., 2017; Cristadoro *et al*., 2018). The most well-known of these are those involved in biological function, such as the typical genome division into coding and non-coding parts or, within genes, the differential conservation shown by distinct codon positions due to the differential evolutionary constraints acting on them (Ikemura, 1985; Sueoka, 1992; Bernardi, 2004). In the present study we intend to capture or approximate the genome information held in these layers using certain metrics (collectively named ‘genome complexity metrics’) to determine whether they show phylogenetic signals and indicate - or not - some kind of evolutionary trend. To do so, we use a group of organisms with a long phylogenetic history: the phylum Cyanobacteria. *SCC* accounts for the global compositional complexity of a DNA sequence encoded by the four nucleotides {A, T, C and G} and shares similarity with McShea’s (1993) operational definition of biological complexity: the degree to which the parts of a morphological structure differ from each other. *SCC_SW_* may account for the complexity due to the partition of the genome into GC-rich and GC-poor segments (as for example, the isochores), which are known to be associated to many functionally relevant properties such as gene density, gene length, retrotransposon density, recombination frequency, etc. (Bernardi *et al*., 1985; Mouchiroud *et al*., 1988; Zoubak *et a*l., 1996; Oliver *et al*., 2004; Bernardi, 2015; Jabbari and Bernardi, 2017). Thus, *SCC_SW_* might capture the genome information gained throughout evolution by the selective forces acting on these important functional elements. On the other hand, *SCC_RY_* accounts for the complexity due to the partition of the genome into segments of different purine/pyrimidine richness. Such strand asymmetries are less directly related to biological function, but this alphabet has been useful to uncover long-range correlations and analyze the evolution of fractal structure in the genomes (Li and Kaneko, 1992; Peng *et al*., 1992; Voss, 1992). Recently, a connection has been found between strand symmetry and the repetitive action of transposable elements during evolution (Cristadoro *et al*., 2018; see also Koonin, 2016a and his concept of ‘fuzzy meaning’ of sequences). The partition given by *SCC_KM_*, to our knowledge, has not been associated to any biological function. Finally, *GS* and *BB* explore the maximum deviation for a given *k*-*mer* between a real and a random genome. *GS* directly compares the observed distribution of *k*-*mer* classes of a real genome with respect to that corresponding to a random one. On the other hand, by calculating the entropy differences between both groups, *BB* measures the relative entropic and anti-entropic fraction of a real genome (Bonnici and Manca, 2016).

From a population genetics perspective, Cyanobacteria can be considered prototypical bacterial species whose populations are evolving under high effective population sizes (Lynch and Connery, 2003), with intermediate mutation rates between those of RNA viruses (higher mutation rate) and lower or higher eukaryotes (lower mutation rates) (Gago *et al*., 2009). Therefore, natural selection is expected to play a major role in the evolution of these organisms. Irrespective of whether mutations (or any source of genetic novelty) are deleterious or beneficial, their destiny will be dictated by the deterministic action of purifying or positive selection, respectively (Lynch, 2007; Koonin, 2016b). This observation is highly pertinent when it comes to appropriately interpreting the phylogenetic signals observed in the metrics of complexity measures and genome parameters following the *in silico* evolutionary processes described by Revell *et al*. (2008). Considering, thus, that selection is a key force in the evolution of Cyanobacteria, most of the *K* values estimated for the metrics may reflect the action of purifying or stabilizing selection, particularly those that are below 1 (all metrics and parameters, except *GS* and %GC). *K* from *GS* is 1, which could be interpreted either as a random drift effect or, more convincingly for this type of organism, as fluctuating selection for a relatively high rate of movement of the optimum (Revell *et al*., 2008). Finally, *K* associated to %GC is much higher than one, which can also be interpreted as the result of an evolutionary process with heterogeneous peak shifts.

Importantly, our study of the evolutionary trends in Cyanobacteria by means of ridge regression found clear differences between metrics of complexity and genome parameters. Four metrics (*SCC, SCC_RY_*, *SCC_SW_* and *GS*) indicate changes toward higher complexity in more evolved clades (long-branch distance with respect to the root of the tree), while *SCC_KM_* does not show any signs of a trend and *BB* shows a negative trend. However, the genome parameters show no evidence of any trend (Figure 2). These results are reinforced when comparatively analyzing trends between metrics and parameters at a lower phylogenetic resolution (Supplementary files 4, 5 and 6). Although metrics used in this work capture different aspects of the evolution of genome complexity in Cyanobacteria (positive trends in *SCC, SCC_RY_* and *GS* vs. negative trends in *SCC_SW_* and *SCC_KM_*), the genome parameters never present any positive trends (Supplementary files 3 and 4). In that respect, although some metrics capture increasing complexity, genome parameters do not. It is interesting to point out the process of selection and genome streamlining of *Synechococcus* and *Prochlorococcus* in clade 97, giving rise to more evolved shorter genomes, which are AT-rich and show a lower number of genes than the rest of Cyanobacteria (Supplementary file 1). As it can be observed, there are statistically significant negative trends in the three genome parameters but also positive trends of *SCC* and *SCC_RY_* metrics (Supplementary files 3 and 4). Therefore, genome reduction in this clade does not imply loss of genome complexity; on the contrary, our study shows that this younger clade also has the most complex genomes of Cyanobacteria, a result in agreement with the high density of functional sequences observed in these free-living organisms (Batut *et al*., 2014).

In summary, considering that selection is a major driver in the evolution of Cyanobacteria, the observed positive trends towards increasing complexity captured by *SCC, SCC_RY_* and *GS* metrics cannot be explained, contrary to Gould (1996), as a passive tendency to increase. The three proofs carried out in order to demonstrate whether positive trends are passive or driven show us that the positive trend is driven and is mainly due to the action of natural selection.

As stated by Day (2012) (see also Corominas-Murtra et al., 2018), a necessary condition for progressive and open-ended evolution is the existence of a parameter (metric) that increases with the evolutionary age of the corresponding organisms. This is what we report here for the case of several metrics, which increase during the evolution of Cyanobacteria.

## Materials and methods

### Phylogenetic Analysis

Ninety-one complete and nearly-complete cyanobacterial genomes were downloaded from GenBank and annotated using Prokka (Seemann, 2014) (see Supplementary file 1). To infer a phylogenomic tree we proceeded first to identify the set of homologous gene families conserved among Cyanobacteria (core genome) using get_homologues.pl pipeline (Contreras-Moreira and Vinuesa, 2013). For this, we used BDBH and OMCL methodologies within get_homologues.pl with the following parameters: a threshold e-value ≤ 10−^10^ for BLAST searches; a minimum percent amino acid identity > 30% between query and subject sequences; and for OMCL, we set the inflation parameter (I) set to 2.0. The consensus core-genome was inferred by the intersection of BDBH and OMCL gene families. To select high-quality phylogenetic markers from the core-genome (i.e. those gene families not showing recombination and/or horizontal gene transfer), we used the software package get_phylomarkers (Vinuesa *et al.,* 2018). By this procedure, we obtained an alignment of 96 top markers. The multiple alignment was cured by eliminating uninformative sites and misaligned positions with Gblocks (Talavera and Castresana, 2007). Finally, a maximum likelihood phylogeny was reconstructed using PhyML (Guindon and Gascuel, 2003) with LG model + I (estimation of invariant sites) + G (gamma distribution) as selected by prottest3 (Darriba *et al.,* 2011). For branch support we used SH statistic within PhyML.

### Genome complexity metrics

*SCC.* Sequence Compositional Complexity of genomes was calculated by using a two-step process. We first obtained the non-overlapping compositional domains comprising the genome sequence, and then applied an entropic complexity measurement able to account for the heterogeneity of such compositional domains. The compositional domains of a given genome sequence are obtained through a segmentation algorithm that was properly designed (Bernaola-Galván *et al*., 1996) by using the Jensen-Shannon entropic divergence (Grosse *et al*., 2002; Lin, 1991) to split the sequence -and iteratively the sub-sequences-into non-overlapping compositional domains which, at a given statistical significance, *s*, are homogeneous and compositionally different from the neighboring domains. Note that this algorithm does not use any scanner window, thus avoiding introducing an artificial parameter. Note also that the statistical significance level *s*, is the probability that the difference between each pair of adjacent domains is not due to statistical fluctuations. By changing this parameter one can obtain the underlying distribution of segment lengths and nucleotide compositions at different levels of detail (Li, 1997), thus fulfilling one of the key requirements for complexity measures (Gell-Mann and Lloyd, 1996). Recent improvements to this segmentation algorithm also allow to segment long-range correlated sequences (Bernaola-Galván *et al*., 2012).

Once a genome sequence was segmented into *n* compositional domains, we computed *SCC* as:

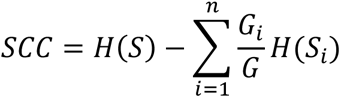

where *S* denotes the whole genomes and *G* its length, *G_i_* the length of the *i* th domain, *S_i_*. *H*(·) = − ∑ *flog*_2_*f* is the Shannon entropy of the distribution of relative frequencies of symbol occurrences {*f*} in the corresponding (sub)sequence (Román-Roldán *et al*., 1998). It should be noted that the above expression is the same one than used in the segmentation process, applying it to the tentative two new subsequences (*n* = 2) to be obtained in each step. Thus, the two parts of the *SCC* computation are based on the same theoretical background.

We apply the above two-step procedure to each of the entire four-symbol cyanobacterial genomes, thus obtaining a *SCC* complexity value for each of them. In addition, we also apply the same procedure to the binary sequences resulting from grouping the four nucleotides into S(C,G) vs. W(A,T) or R(A,G) vs. Y (T,C), or K(A,C) vs. M(T,G), then obtaining *SCC_SW_*, *SCC_RY_* and *SCC_KM_* metrics, respectively. These three additional metrics are partial complexities providing complementary views of genome complexity to that obtained with the four-symbol sequence (Bernaola-Galván *et al*., 1999; Bernaola-Galván *et al*., 2004).

*GS.* The Chaos Game Representation (*CGR*, Jeffrey, 1990; Almeida *et al*., 2001) is an image derived from a genome where each point of the image corresponds to a given *k*-*mer* level of analysis. If the genome sequence is a random collection of bases, the *CGR* will be a uniformly filled square image. On the bases of building a *CGR* for a particular genome, we define a corresponding genomic signature (*GS*) that is a numerical value obtained for a particular *k*-*mer* level by comparing point-by-point the difference between the *CGR*’s of a real genome and a random genome of the same length. In order to make it comparable, the pixel values of the images are normalized. As stated, the size of the images generated depends on the *k*-*mer* used. For a given *k*-*mer*, we have *4^k^* different words and the corresponding image *4^k^* pixels too. To build a frequency table for each *k*-*mer* minus the expected frequency for a random genome is equivalent to the difference between the *CGR* images of a real and a random genome. In fact, if *G* is the size of the genome to analyze, the expected value (*EV*) for a given *k*-*mer* is given by *EV=G/(4^k^)*. This value is used to normalize to 1 the values of the *k*-*mers* obtained for each of the genomes analyzed. We then define the *GS* as:

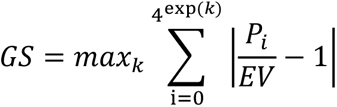

where *P_i_* is the relative frequency of the *k*-*mer i*.

*BB.* The biobit is a metric of genome complexity that is derived from the comparison between the *k*-*mer* that yields the maximum entropy of a given random genome and the corresponding entropy of the real genome of the same length (Bonnici and Manca, 2016). The authors demonstrated that the entropy of a real genome of length *G, E_2L(G)_* takes a value between the maximum (*2log_4_(G)* or *2L(G)*) and the minimum (*L(G)*) entropy. On the other hand, Bonnici and Manca (2016) define and measure two additional components, that they call entropic (*E(G*)) and anti-entropic (*A(G)*) of a real genome, in such a way that *A(G) + E(G) = L(G)*. Then, the entropy of those components are given by *E(G) = E_2L(G)_ - 2L(G)* and *A(G) = 2L(G) - E_2L(G)_*, respectively.

Finally, the biobit *BB* of a genome (*BB(G)*), is a non-linear combination of the two entropic and anti-entropic components given by:

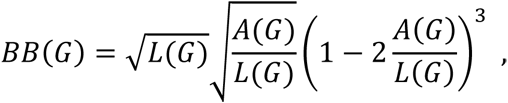

where 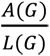 is the anti-entropic fraction of the genome and 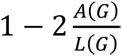 is the corresponding entropic fraction. Both components vary between 0 and 1.

#### Standard genome parameters

Finally, we have also included three standard genome parameters: genome size, %*GC* and number of genes.

### Phylogenetic signal

We used the phylogenetic tree of Cyanobacteria to test the existence of a phylogenetic signal in the genome complexity metrics and genome parameters through Blomberg *et al*. (2003) *K*-statistic in the picante package for *R* (Kembel *et al*., 2010). *K* ranges from 0 to ∞. *K* values significantly higher than zero are indicative of the presence of a phylogenetic signal or, in other words, that closely related species resemble more in the study trait than expected by chance. *K* = 1 is the value expected under Brownian evolution.

### Phylogenetic correlations

We have examined the correlation between genome parameters and metrics of genome complexity after correcting the phylogenetic signal. Pearson *r* value between variables was computed as the phylogenetic trait variance-covariance matrix between two variables and significance tested against a *t*-distribution with *n-2* degrees of freedom. We used the R code provided by Liam Revell to perform Pearson correlation with phylogenetic data (http://blog.phytools.org/2017/08/pearson-correlation-with-phylogenetic.html). The *p*-value obtained with this procedure is the same as that provided by a phylogenetic generalized linear square model. As we run multiple phylogenetic correlations, we corrected *p*-values by false discovery rates.

### Evolutionary trends

We tested the existence of an evolutionary trend in the genomic complexity measures and genome parameters by fitting a ridge regression of each of these genomic values against evolutionary age (i.e., tip-to-root or node-to-root distances). Significance was tested against 10,000 slopes obtained under Brownian motion simulations with the help of the *search.trend* function in the RRphylo package for R (Castiglione *et al.,* 2019).

## Acknowledgements

This work was supported by grants from the Spanish Minister of Science, Innovation and Universities (former Spanish Minister of Economy and Competitiveness) to AM (project SAF2015-65878-R), JLO (project AGL2017-88702-C2-2-R) and AL (project PGC2018-099344-B-I00), grant from Generalitat Valenciana to AM (project Prometeo/2018/A/133), and co-financed by the European Regional Development Fund (ERDF). This project was also supported by a Fulbright fellowship (Spanish Minister of Science, Innovation and Universities) to AM for a sabbatical leave at Harvard University.

## Additional files Supplementary files

**Supplementary file 1. Table S1.**
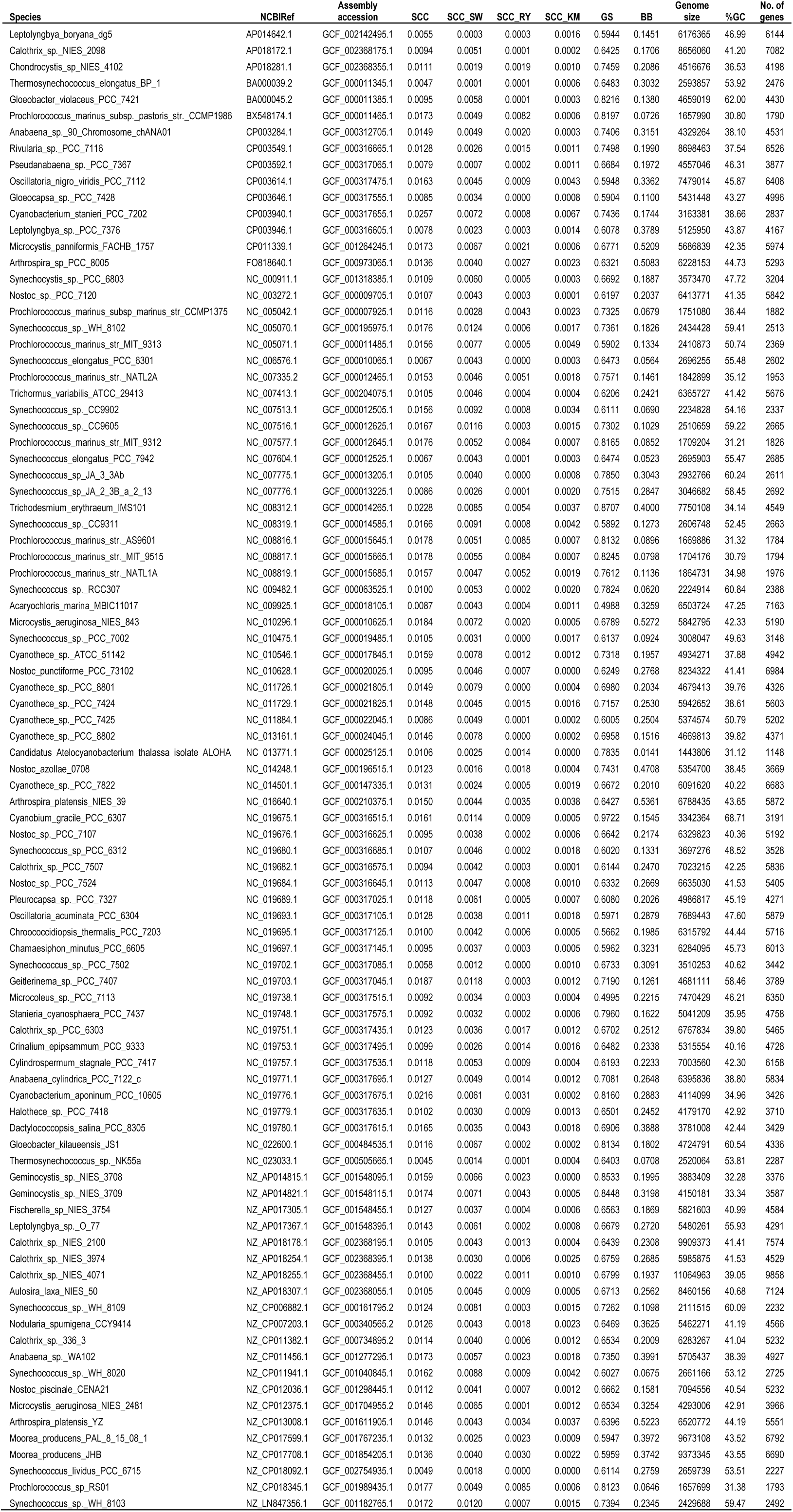
General genome features, genome parameters, and metrics of genome complexity of Cyanobacteria genomes

**Supplementary file 2. Figure S1.**
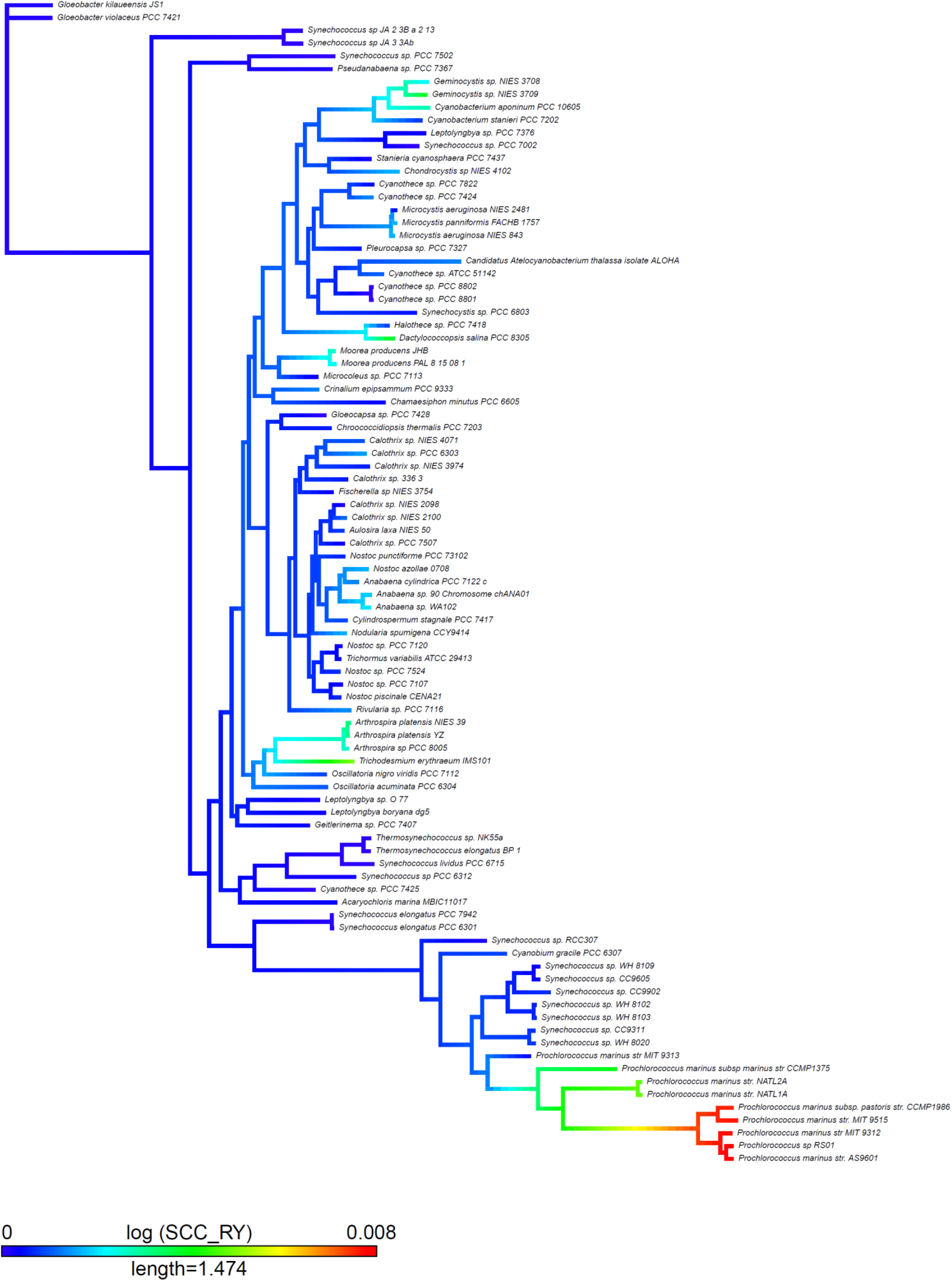
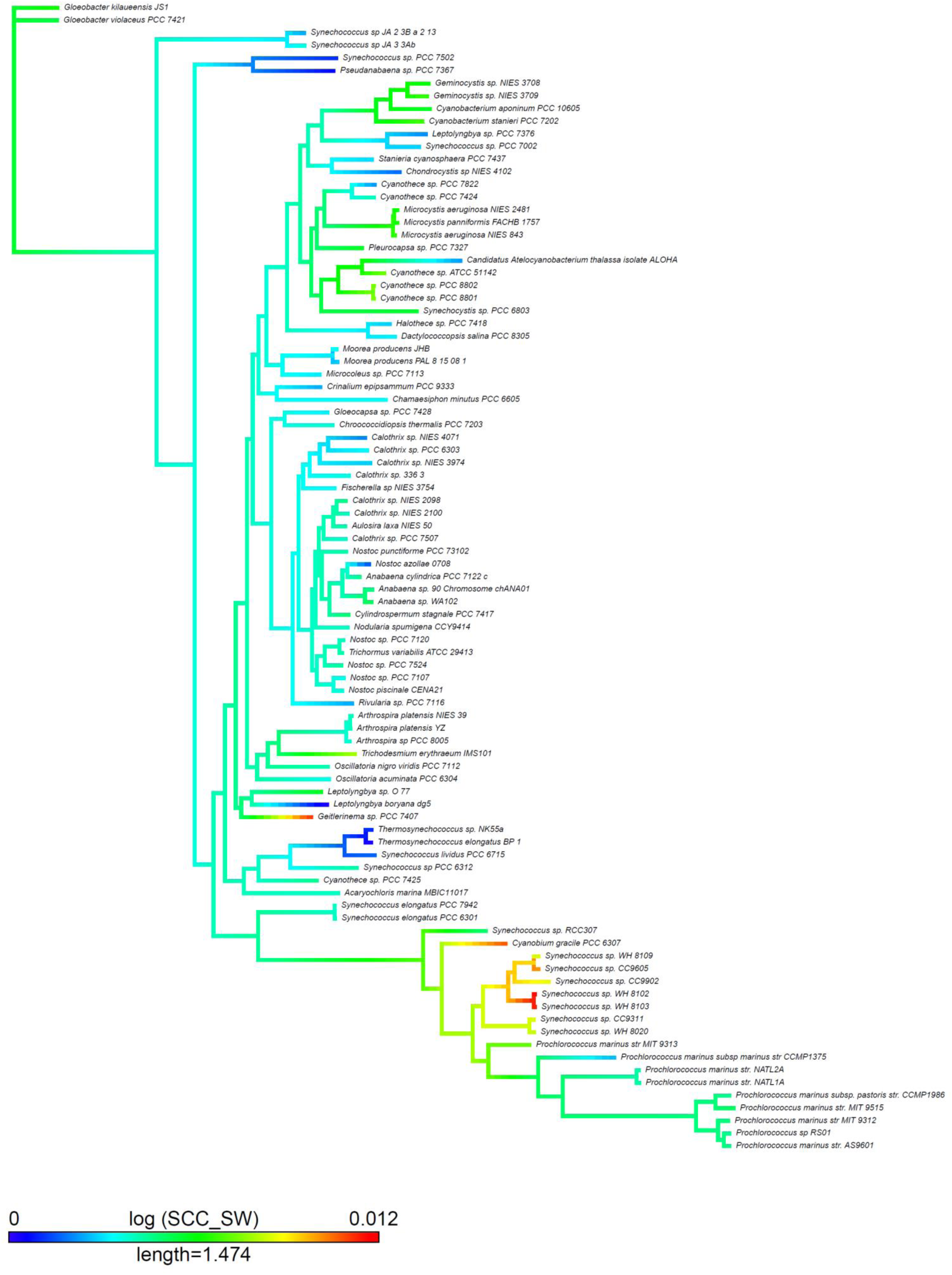
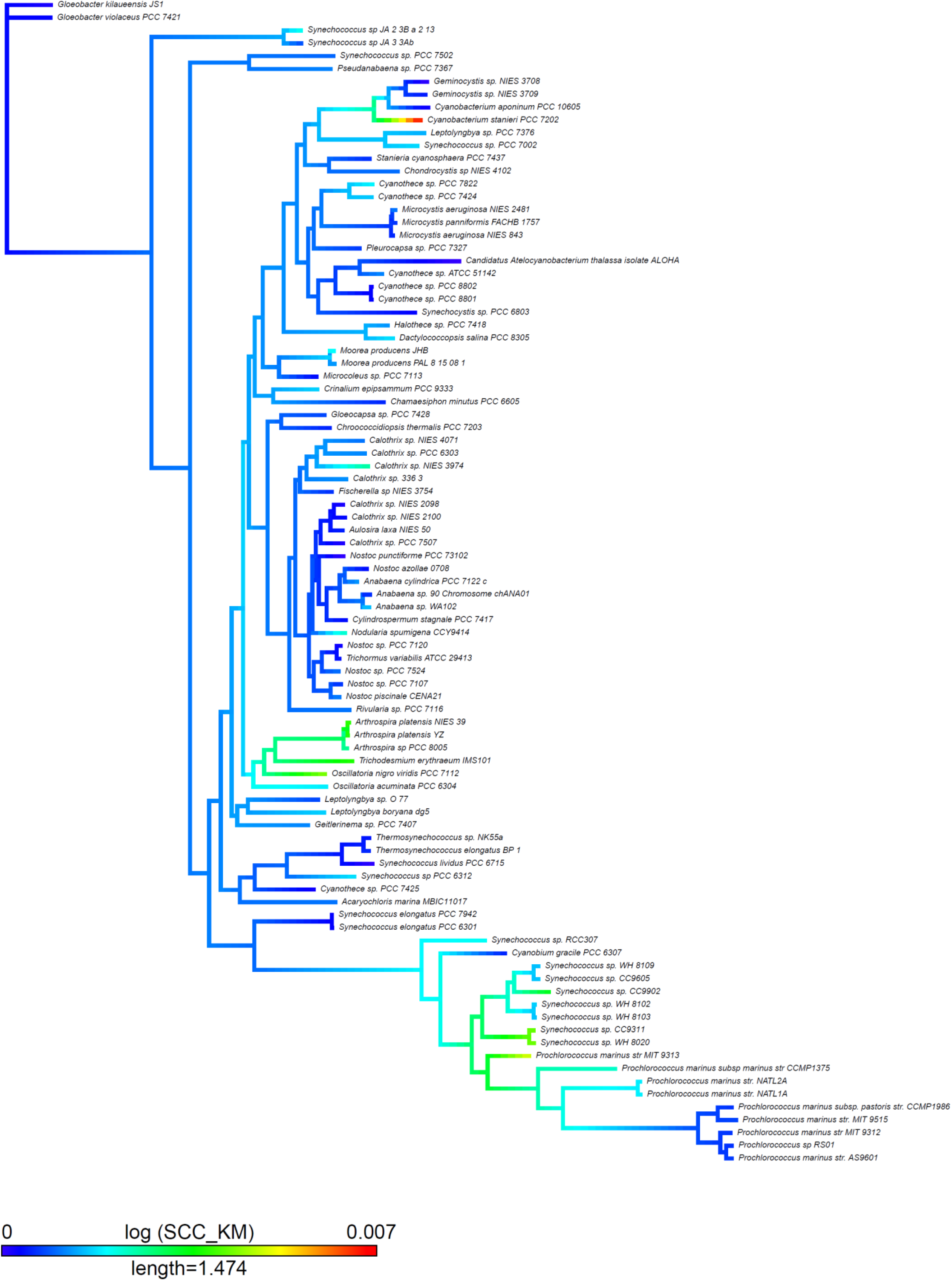
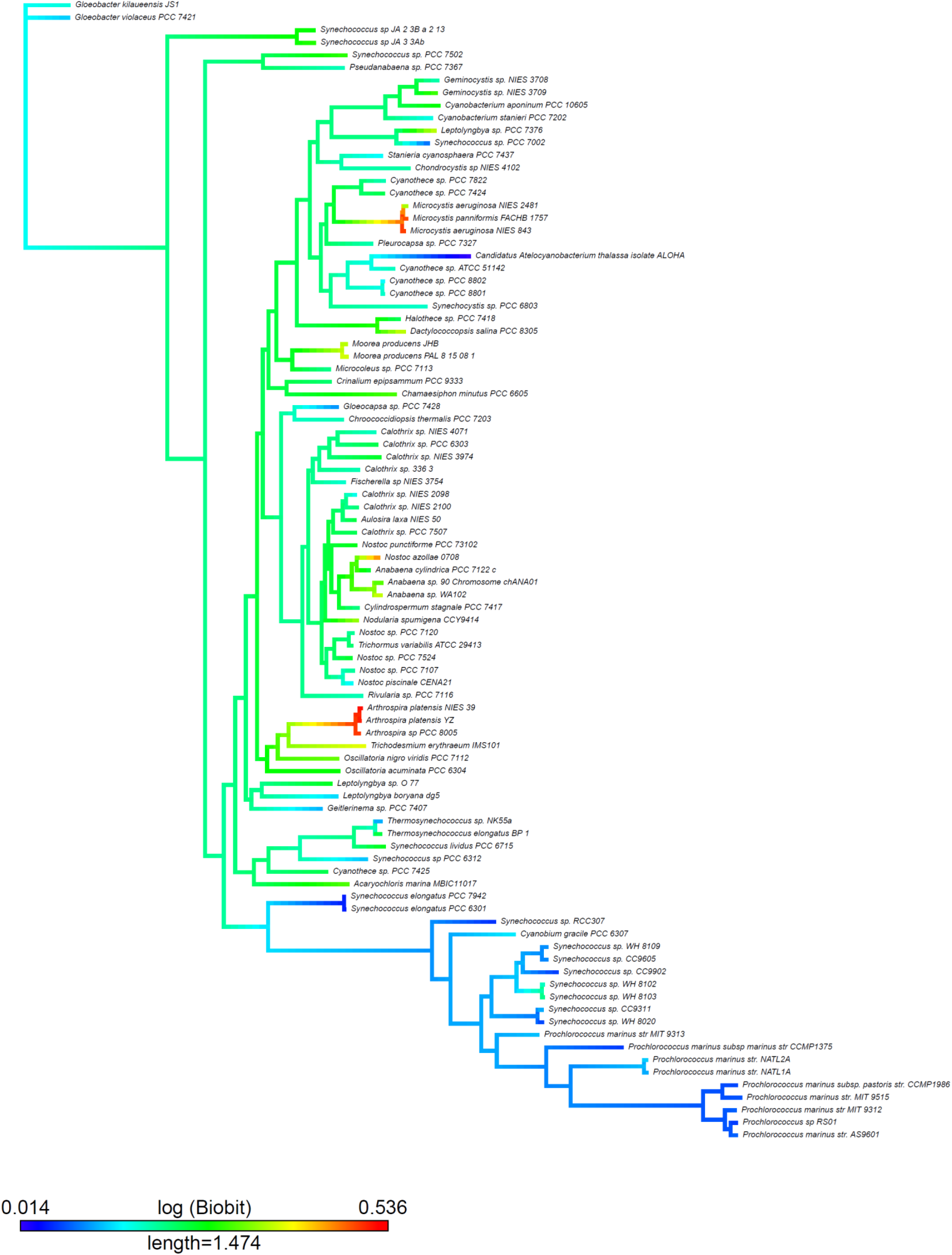
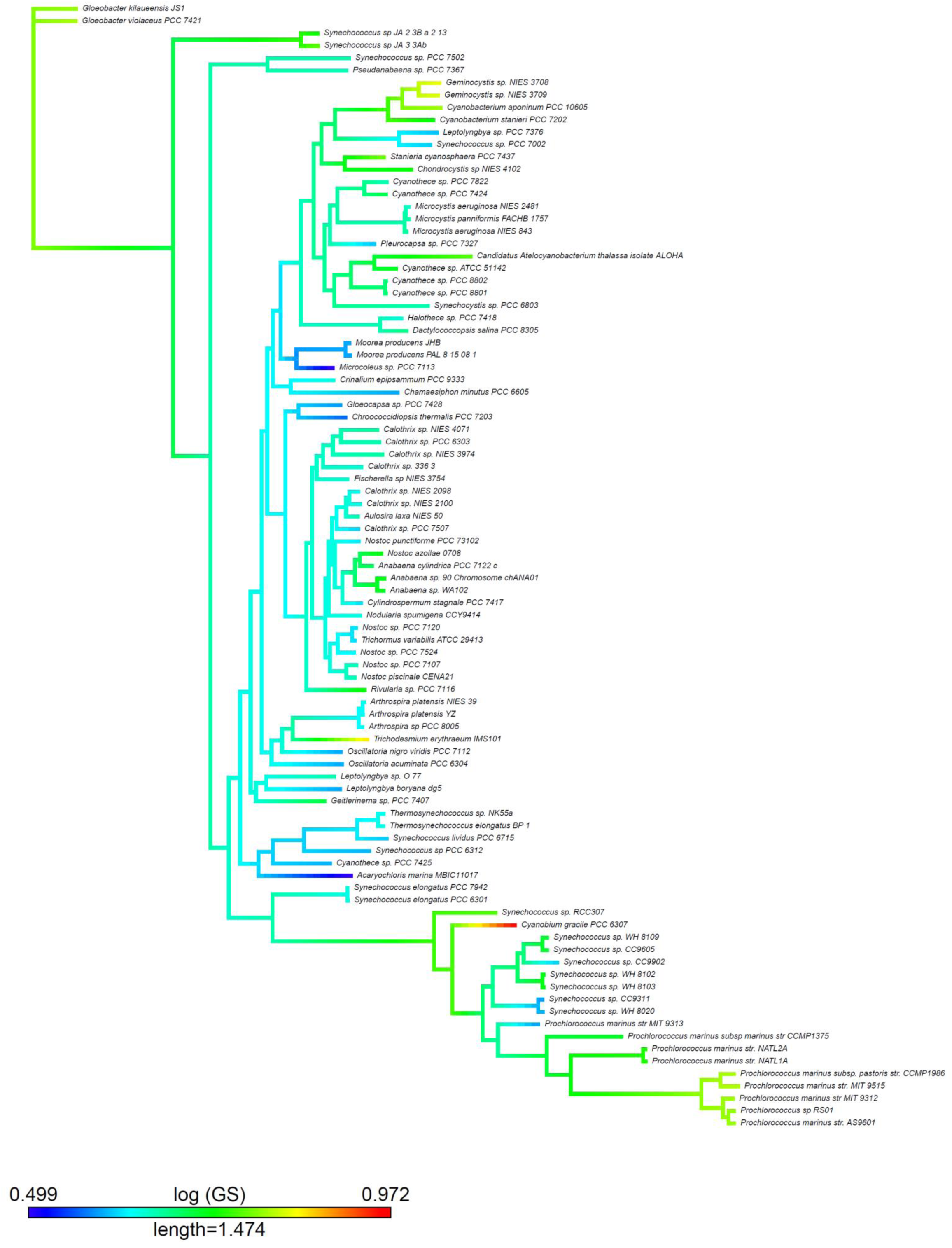
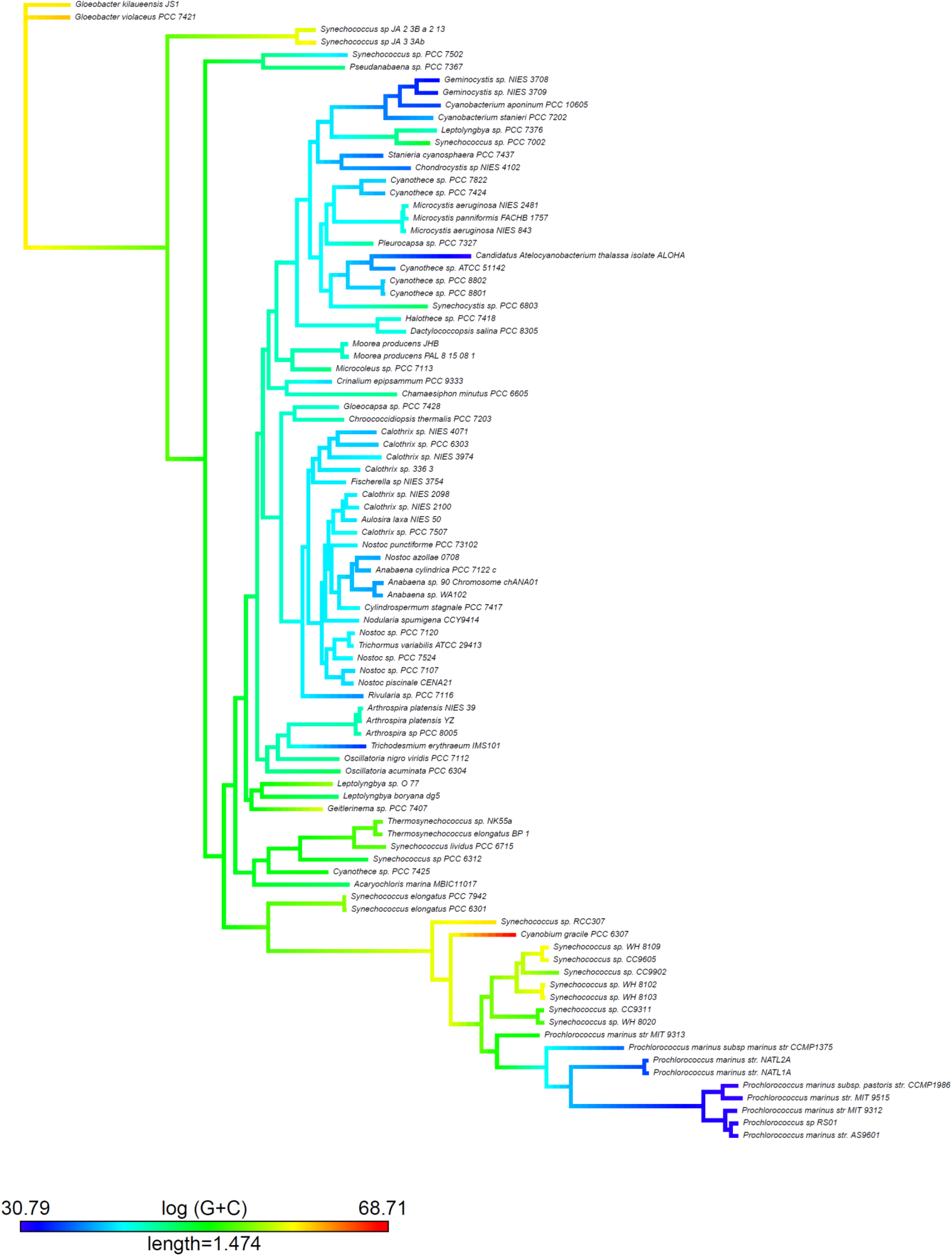
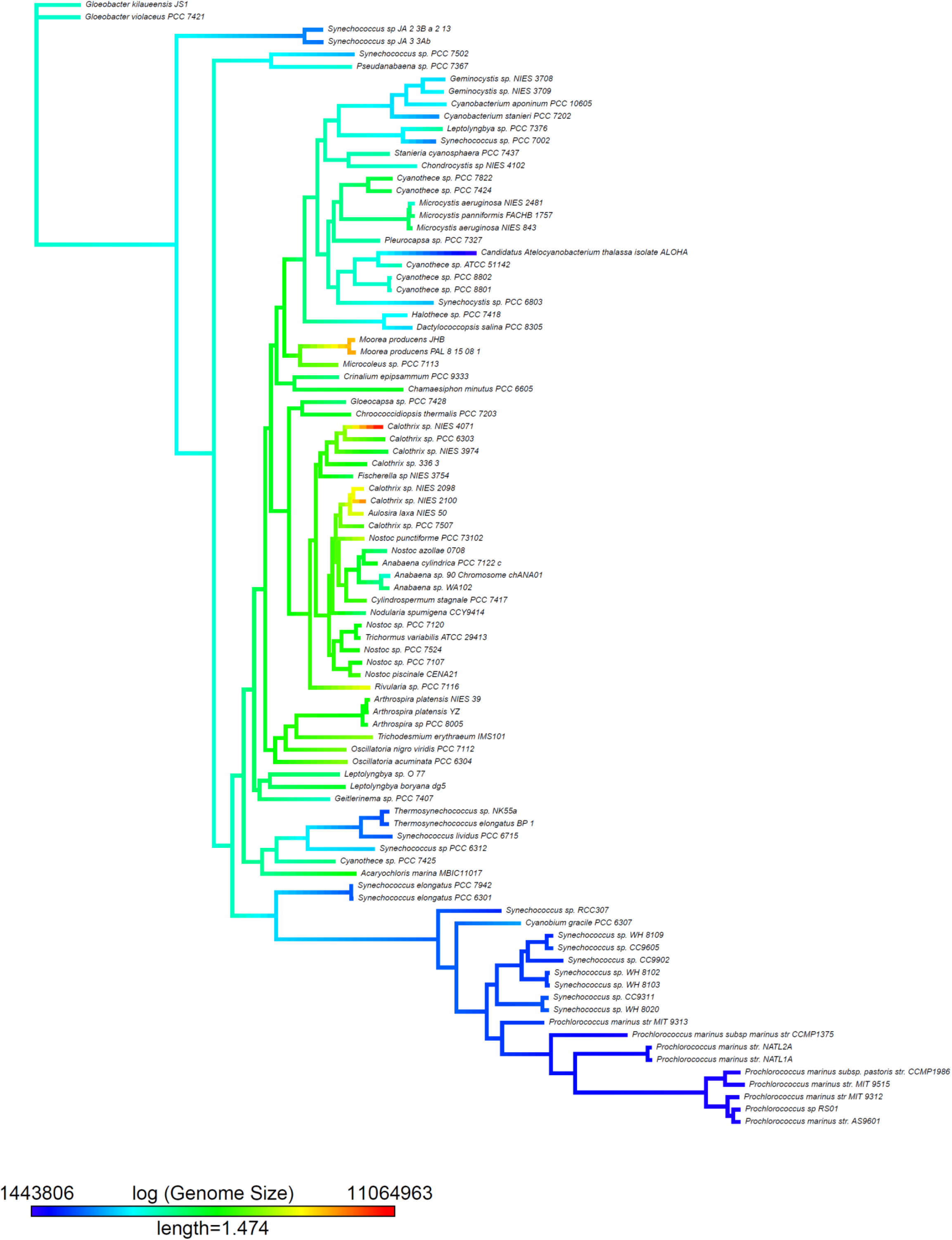
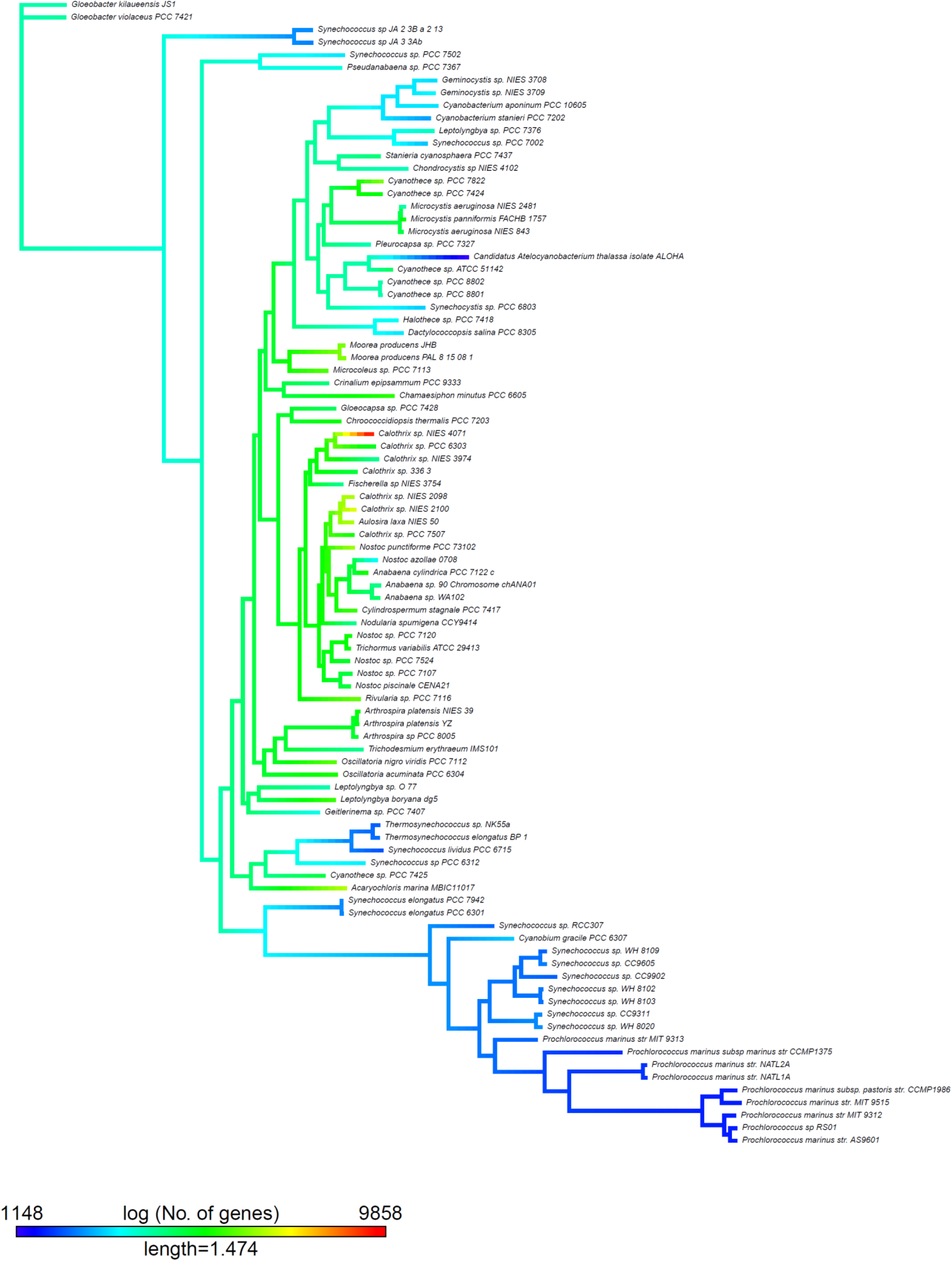
Cyanobacteria tree mapping the rest of the metrics and parameters. We used two functions (*contMap* and *fastAnc*) from the *phytools* R package (Revell, 2012). The *contMap* R function allows plotting a tree with a mapped continuous character, such as any of our complexity measures. The mapping is accomplished by estimating states at internal nodes using maximum likelihood with the function *fastAnc* and interpolating the states along each edge using equation 2 of Felsenstein (1985).

**Supplementary file 3. Figure S2.**
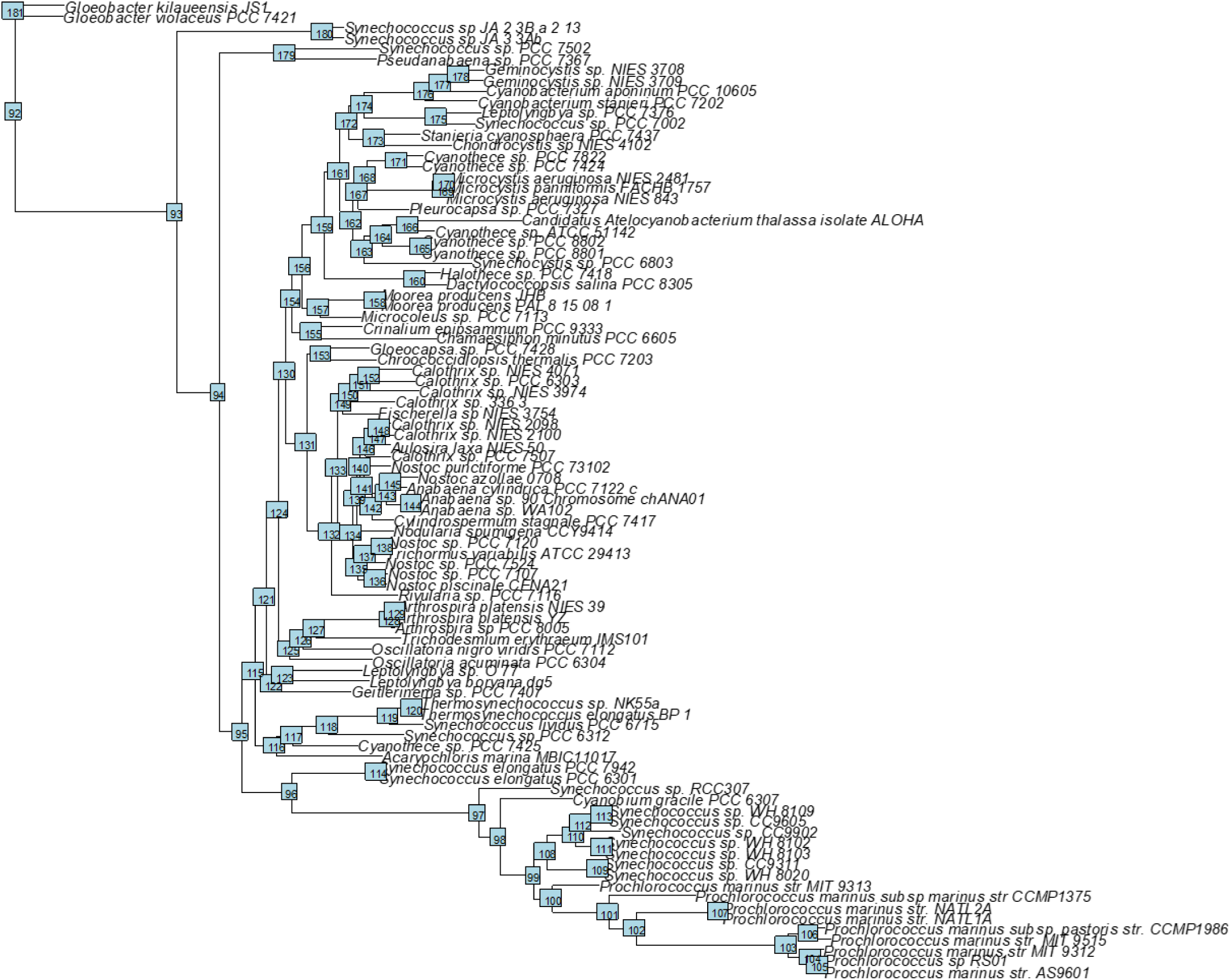
Indication on the phylogenetic tree of the Cyanobacteria the location of the four monophyletic clades (97, 132, 162 and 172) where evolutionary trends of metrics and genome parameters were evaluated.

**Supplementary file 4. Table S2.**
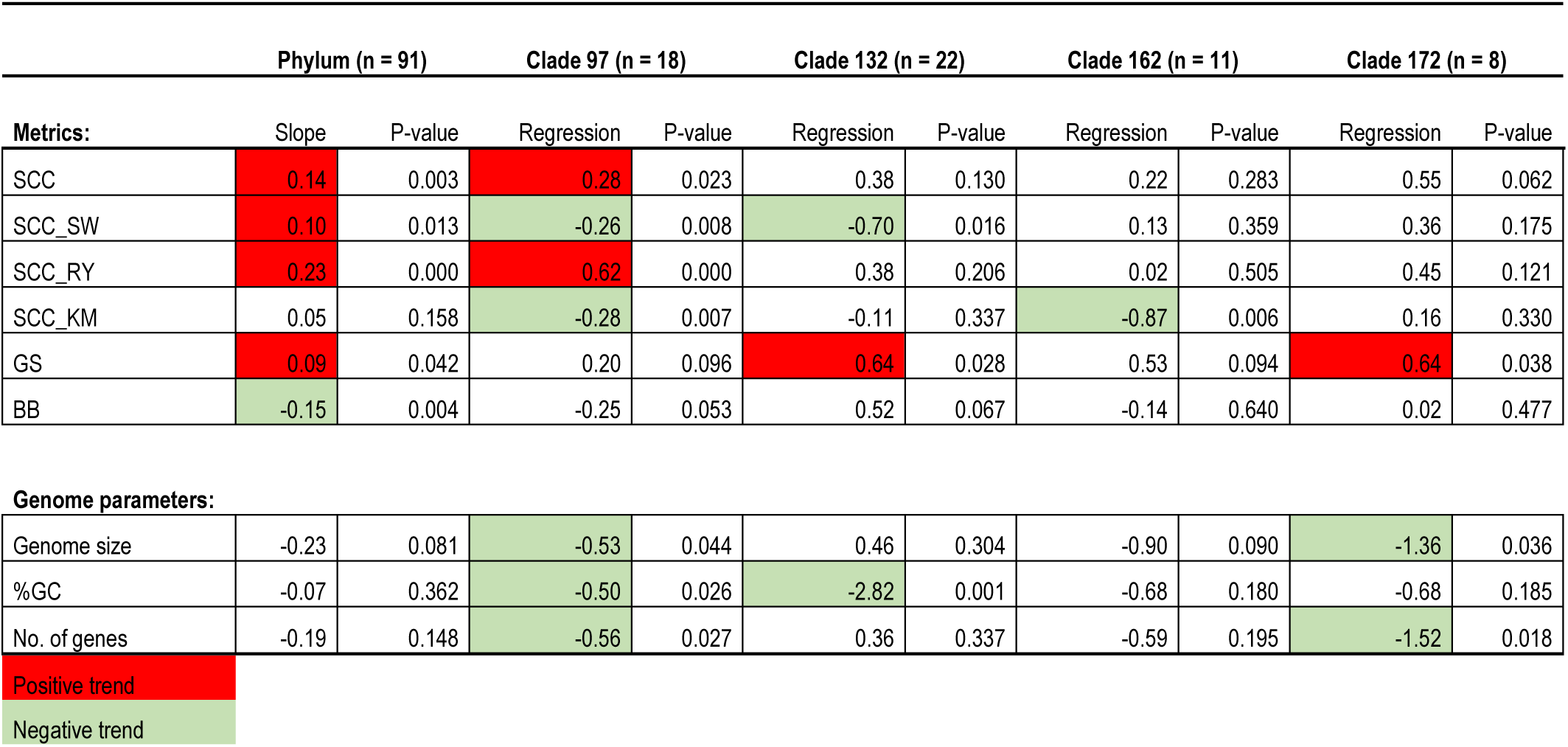
Ridge regression of genome complexity metrics and genome parameters versus age (distance from the root) in the phylum and the four selected monophyletic clades.

**Supplementary file 5. Figure S3.**
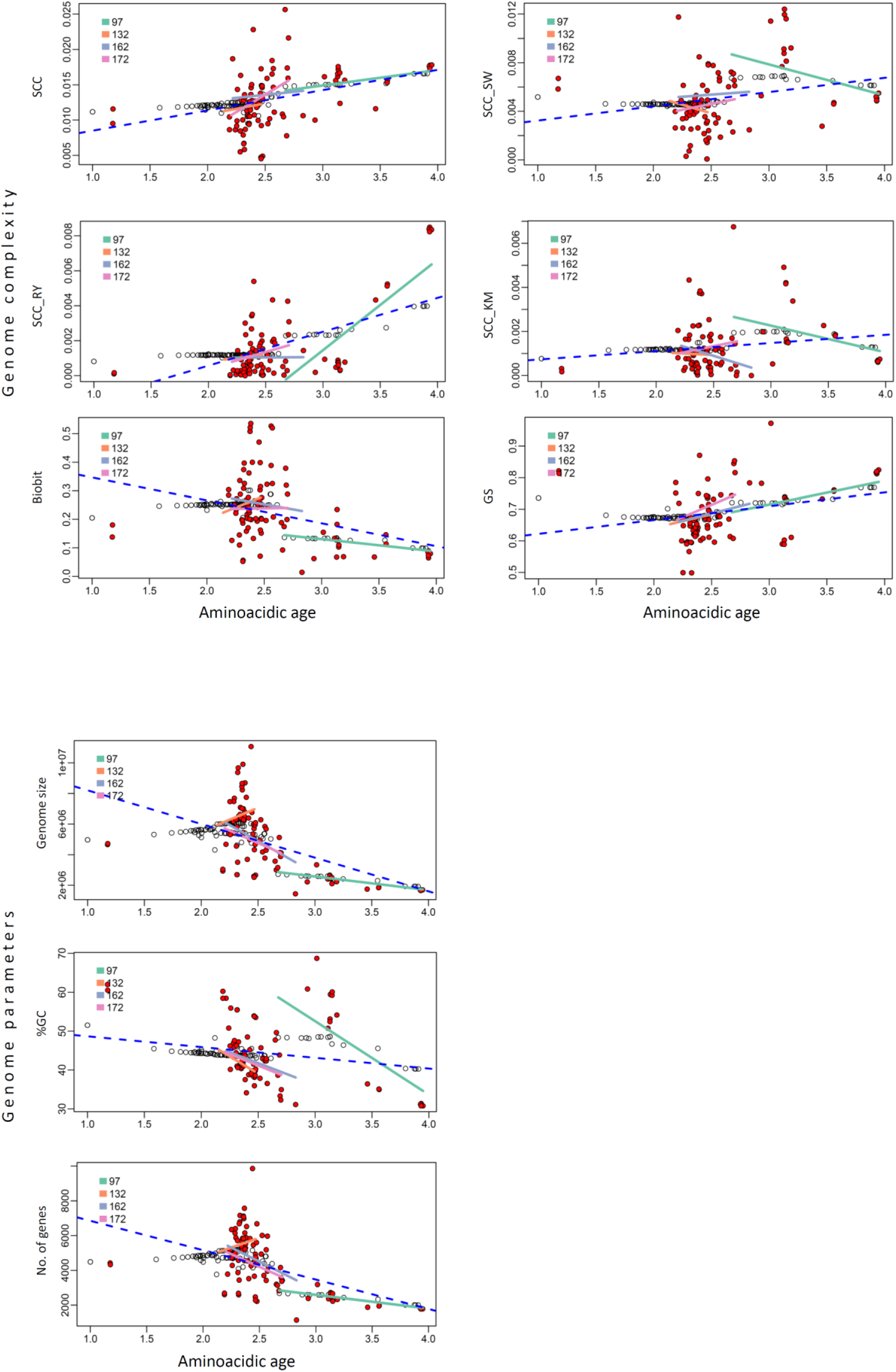
Phylogenetic trends of genomic complexity metrics and standard genome parameters in the clades 97, 132, 162 and 172. The estimated genomic value for each tip (red dots) or node (black dots) in the phylogenetic tree is regressed against its evolutionary age (i.e. distance from the root). The shaded area of the plot shows a 95% confidence interval of the estimated genomic values. The statistical significance of the regression is tested against 10,000 slopes obtained under simulated Brownian evolution.

**Supplementary file 6. Table S3.**
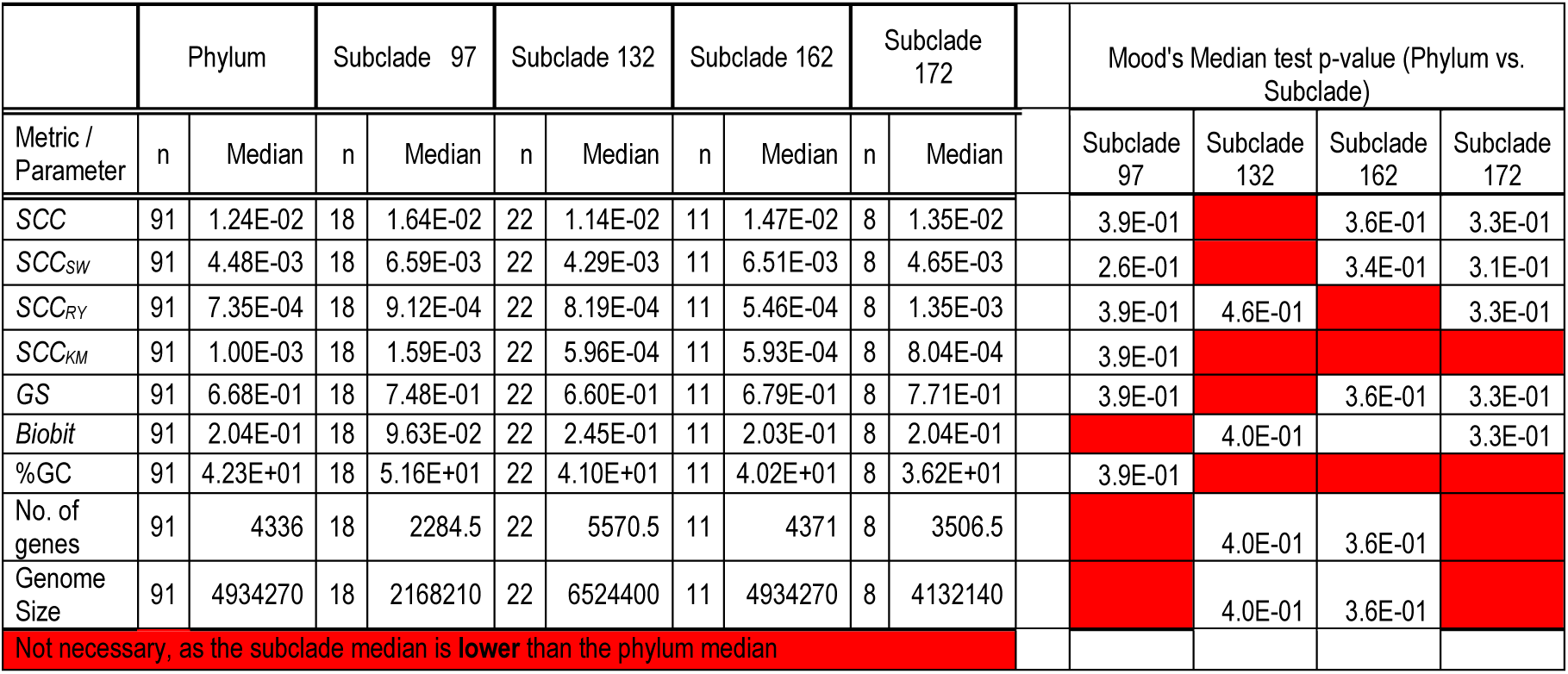
Sub-clade proof, second sub-clade test. Median values in the entire phylum and the four sub-clades. In the protocol adopted here, a subclade drawn from the tail is defined as a monophyletic subset chosen such that the mean fits distribution is greater than the mean of the parent distribution (McShea, 1994).

|  | Phylum |  | Subclade 97 |  | Subclade 132 |  | Subclade 162 |  | Subclade 172 |  | Mood's Median test p-value (Phylum vs. Subclade) |  |  |  |
| --- | --- | --- | --- | --- | --- | --- | --- | --- | --- | --- | --- | --- | --- | --- |
| Metric / Parameter | n | Median | n | Median | n | Median | n | Median | n | Median | Subclade 97 | Subclade 132 | Subclade 162 | Subclade 172 |
| SCC | 91 | 1.24E-02 | 18 | 1.64E-02 | 22 | 1.14E-02 | 11 | 1.47E-02 | 8 | 1.35E-02 | 3.9E-01 |  | 3.6E-01 | 3.3E-01 |
| SCC<sub>SW</sub> | 91 | 4.48E-03 | 18 | 6.59E-03 | 22 | 4.29E-03 | 11 | 6.51E-03 | 8 | 4.65E-03 | 2.6E-01 |  | 3.4E-01 | 3.1E-01 |
| SCC<sub>RY</sub> | 91 | 7.35E-04 | 18 | 9.12E-04 | 22 | 8.19E-04 | 11 | 5.46E-04 | 8 | 1.35E-03 | 3.9E-01 | 4.6E-01 |  | 3.3E-01 |
| SCC<sub>KM</sub> | 91 | 1.00E-03 | 18 | 1.59E-03 | 22 | 5.96E-04 | 11 | 5.93E-04 | 8 | 8.04E-04 | 3.9E-01 |  |  |  |
| GS | 91 | 6.68E-01 | 18 | 7.48E-01 | 22 | 6.60E-01 | 11 | 6.79E-01 | 8 | 7.71E-01 | 3.9E-01 |  | 3.6E-01 | 3.3E-01 |
| Biobit | 91 | 2.04E-01 | 18 | 9.63E-02 | 22 | 2.45E-01 | 11 | 2.03E-01 | 8 | 2.04E-01 |  | 4.0E-01 |  | 3.3E-01 |
| %GC | 91 | 4.23E+01 | 18 | 5.16E+01 | 22 | 4.10E+01 | 11 | 4.02E+01 | 8 | 3.62E+01 | 3.9E-01 |  |  |  |
| No. of genes | 91 | 4336 | 18 | 2284.5 | 22 | 5570.5 | 11 | 4371 | 8 | 3506.5 |  | 4.0E-01 | 3.6E-01 |  |
| Genome Size | 91 | 4934270 | 18 | 2168210 | 22 | 6524400 | 11 | 4934270 | 8 | 4132140 |  | 4.0E-01 | 3.6E-01 |  |
| Not necessary, as the subclade median is lower than the phylum median |  |  |  |  |  |  |  |  |  |  |  |  |  |  |

